# Genome dilution by cell growth drives starvation-like proteome remodeling in mammalian and yeast cells

**DOI:** 10.1101/2023.10.16.562558

**Authors:** Michael C. Lanz, Shuyuan Zhang, Matthew P. Swaffer, Inbal Ziv, Luisa Hernández Götz, Frank McCarty, Daniel F. Jarosz, Joshua E. Elias, Jan M. Skotheim

**Affiliations:** Department of Biology, Stanford University, Stanford CA 94305; Department of Chemical and Systems Biology, Stanford University, Stanford CA 94305; Chan Zuckerberg Biohub San Francisco, Stanford University, Stanford CA 94305

## Abstract

Cell size is tightly controlled in healthy tissues and single-celled organisms, but it remains unclear how size influences cell physiology. Increasing cell size was recently shown to remodel the proteomes of cultured human cells, demonstrating that large and small cells of the same type can be biochemically different. Here, we corroborate these results in mouse hepatocytes and extend our analysis using yeast. We find that size-dependent proteome changes are highly conserved and mostly independent of metabolic state. As eukaryotic cells grow larger, the dilution of the genome elicits a starvation-like proteome phenotype, suggesting that growth in large cells is limited by the genome in a manner analogous to a limiting nutrient. We also find that this phenomenon explains many proteomic changes ascribed to yeast aging. Overall, our data suggest that genome concentration is a universal determinant of proteome content in growing cells.

## INTRODUCTION

Cell size is intimately related to cell function and therefore tightly controlled. Macrophages must be large enough to engulf pathogens, whereas lymphocytes must be small enough to squeeze through tight passages. However, apart from such exceptional cases, it is unclear why most cell types adopt their characteristic size. Nevertheless, we know that cell size is tightly controlled in both human tissues and single-celled organisms, suggesting universal importance. For example, budding yeast and mammalian epithelial cells correct for variations birth size by adjusting cell growth in their G1 phases ^1,2^. In fission yeast such size regulation takes place in G2 ^3^. Each mechanism ensures that cells that were born large or small finish the division cycle at a similar target size.

To control cell size, cells must be able to “sense” how large they are. Recent mechanistic investigations of size control have focused on individual proteins whose concentrations reflect cell size. Several cell division inhibitors were found to be progressively diluted by cell growth, which ultimately promotes cell division (reviewed in reference ^4^). For example, the G1/S transcriptional inhibitors Whi5 in budding yeast and Rb in human cells are diluted in G1 ^5–7^. In *Arabidopsis*, the dilution of the cell cycle kinase inhibitor Krp4 regulates cell size in the shoot stem cell niche ^8^, and in *Chlamydomonas* (green algae), the dilution of the RNA-binding protein TNY1 regulates the expression of a cell cycle transcripts ^9^. Conversely, cell division activators may behave in the opposite way^10^. For example, the fission yeast cell cycle phosphatase Cdc25, which promotes mitosis, becomes increasingly concentrated in large cells ^11^. Thus, the differential size-scaling of cell cycle regulatory proteins can regulate the division cycle to maintain a desired target size range.

Despite the ubiquity of size regulation in cell biology, it had been unclear how natural variations in cell size would generally affect cell physiology. Apart from the recent discovery that several cell cycle regulatory proteins change concentration with cell size, it was thought that the concentrations of most individual proteins and mRNAs were kept constant as cells grow ^12–18^. However, recent mass spectrometry studies of cultured human cells have potentially revealed a more general link between cell size and cell physiology ^19,20^. These studies showed that the concentrations of many individual proteins continuously change as cells grow larger, despite the fact that total mRNA ^18,21,22^ and protein ^12,17,21,23–25^ concentrations remain roughly constant. Proteins whose concentration increases or decreases with cell size are termed super- and sub-scaling, respectively (**Figure 1A**). Each protein’s degree of size-scaling can be quantified by regressing its relative change in concentration against the relative change in cell size, producing a slope value (**Figure 1B**). A slope value of 1 corresponds to a doubling of the concentration with a doubling cell size, a value of 0 corresponds to a constant concentration, and a value of -1 corresponds to halving the concentration with a doubling of cell size (*i.e.*, dilution) (**Figure 1B**). The revelation that individual proteins adopt a broad distribution of size-scaling behaviors could explain how cell size generally influences cell physiology. Namely, the deviation from perfect size scaling provides a means for large and small cells of the same type to be biochemically different.

**Figure 1:**
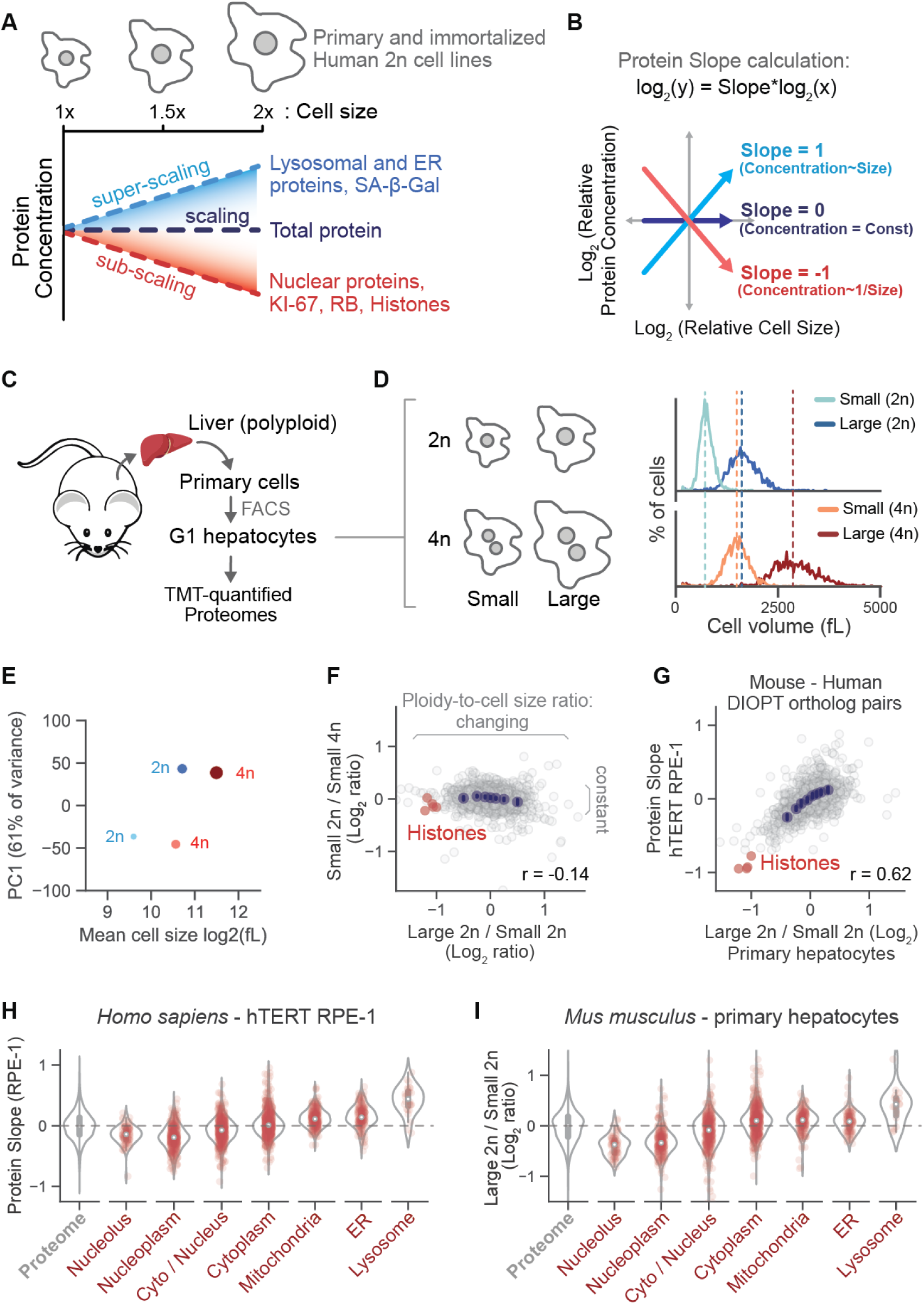
Ploidy-to-cell size ratio, rather than cell size *per se*, drives conserved changes in the mammalian proteome. **(A)** Schematic summary of the scaling relationships between protein concentration and cell size as described previously in cultured human cell lines ^19^. Schematic adapted from Lanz *et al.*^19^ **(B)** Derivation of the Protein Slope value. Protein Slopes describe the size-scaling behavior of each individual protein. Proteins with a slope of 0 maintain a constant cellular concentration regardless of cell volume. A slope value of 1 corresponds to an increase in concentration that is proportional to the increase in volume and a slope of -1 corresponds to dilution (1/volume). Schematic adapted from Lanz *et al.*^19^ **(C)** Schematic illustrating TMT-based proteomic analysis of mouse primary hepatocytes of different size and ploidy. **(D)** Small and large hepatocytes with 2n or 4n genome content were isolated using FACS. G1 phase populations were attained using a fluorescent FUCCI reporter (Figure S1). Cell size distributions were determined using a Coulter counter. **(E)** Principal component analysis of the relative concentration of each protein in small or large G1 cells of either 2n or 4n ploidy. The 1st principal component is plotted against cell volume. Dot size represents mean cell volume. **(F)** Correlation of concentration changes for individual proteins. 2,096 proteins are plotted. Only proteins with at least 5 independent measurements (see methods) are shown (see Table S1 for all proteins). Log_2_ ratios are calculated from the relative change in a protein’s concentration between small and large cells. The log_2_ (large / small) concentration ratio is proportional to the protein slope value when comparing cells with a 2-fold difference in cell size. **(G)** Correlation of the size-scaling behaviors of orthologous proteins in mouse and human cells (data from *Lanz et al.* ^19^). The log_2_ (large / small) concentration ratio is proportional to the protein slope value because the large hepatocytes are roughly 2-fold larger than the small hepatocytes. **(H and I)** Scaling behavior of mouse and human proteins strictly parsed by subcellular localization (see methods). Human data is re-plotted from Lanz *et al.* ^19^ The log_2_ (large / small) concentration ratio is directly proportional to the protein slope value because the large hepatocytes are roughly 2-fold larger than the small cells.

That the proteome continuously remodels as cultured human cells grow larger raises three important questions. First, what is the mechanism by which increasing cell size remodels proteome content? Second, how conserved is this phenomenon? Third, how do these proteome changes ultimately reflect changes in cell physiology? Here, we demonstrate that changes genome concentration (*i.e.*, ploidy-to-cell size ratio), rather than cell size *per se*, remodel proteome content in yeast and mammalian cells. These proteome changes are highly conserved and mostly independent of metabolic state. In both yeast and human cells, the dilution of the genome elicits a starvation-like proteome phenotype, suggesting that growth in large cells is limited by the genome in a manner analogous to a limiting nutrient. Overall, we conclude that genome dilution is likely a universal determinant of proteome content for growing cells and that the proteomes of aged or senescent cells may be primarily determined by their large size.

## RESULTS

### The ploidy-to-cell size ratio determines proteome content in diploid and tetraploid mouse hepatocytes

To determine whether the size-dependent proteome remodeling is conserved across mammalian cell types, we measured how cell size and ploidy impacted the proteomes of primary mouse hepatocytes. Mouse hepatocytes were chosen because their ploidy in the mouse liver is heterogeneous, so we could simultaneously test the effects of cell size and ploidy in a single experiment. We derived primary cell lines containing both diploid and tetraploid hepatocytes (**Figure 1C**). We then measured the proteomes of FACS-isolated small and large G1 cells that were either diploid or tetraploid (**Figure 1D and S1**). Small 4n (dark blue) and large 2n (dark red) populations were approximately the same average size, but their proteome content was different (**Figure 1E,F; Table S1**). In contrast, the proteome content of small 2n and small 4n cells was similar despite a 2-fold difference in size (**Figure 1E,F**). Importantly, small 2n and small 4n cells have a similar cell size-to-ploidy ratio. Consistent with what was previously found in human cell lines, the relative concentration of mouse histone proteins also reflected the relative concentration of the genome (**Figure 1F**). These results demonstrate that ploidy-to-cell size ratio determines proteome content in cultured mammalian cells and suggest that surface area-to-volume ratio is largely irrelevant. This finding is consistent with a previous experiment where mammalian cells were chemically manipulated to endoreduplicate using an aurora kinase inhibitor ^19,26^. In these experiments, proteome content also primarily changed with the ploidy-to-cell size ratio rather than cell size *per se*.

After establishing that the ploidy-to-cell size ratio determined proteome content in primary mouse hepatocytes, we next tested if the same proteins were similarly affected by this ratio in mouse and human cells. To do this, we directly compared the size-scaling behavior of individual mouse and human proteins by matching them based on sequence orthology (DIPOT) ^27^. Size-dependent changes to the mouse and human proteome were highly correlated (**Figure 1G, Table S2**). Moreover, differential size-scaling in organelle protein content, a prominent signature of cell size ^19^, was the same in mice and human cells (**Figure 1H,I**). The similarity of size-dependent proteome changes in these distinct cell types suggests that the ploidy-to-cell size ratio determines the proteome content of most proliferating mammalian cells.

### Conservation of proteome size-scaling from mammals to yeast

To test if size-dependent proteome remodeling is conserved across eukaryotes, we turned to the model single-celled organism budding yeast (*S. cerevisiae*). We chose to examine budding yeast because it does not have the drawbacks associated with cultured mammalian cells, which are grown using an *in vitro* regimen that clearly differs from how mammalian cells grow and proliferate *in vivo*. Thus, we sought to investigate the relationship between cell size and the proteome using a more physiological model system.

To measure how the proteome changes with budding yeast cell size, we generated different-sized yeast cells using two orthogonal approaches and then measured proteome content using a label-swapping triple-SILAC workflow described previously ^19^ (**Figure S2**). First, we used mutant strains that have an average size that is either smaller or larger than a wildtype strain (“Size mutants”; **Figure 2A and S3**) ^28^. Second, we used a strain that conditionally expresses a single G1 cyclin (*CLN1*) under the control of a β-estradiol responsive promoter. In the absence of β-estradiol, these cells arrest in G1 because they lack expression of any of the three G1 cyclins (*cln2Δcln3Δ*) ^29^. Unlike yeast arrested by mating pheromone, biomass accumulates after washout at a rate that is initially near-exponential (**Figure 2B**). By sampling cultures that have been arrested for different lengths of time, we isolated cells of different sizes (“G1 arrest time”, **Figure 2A and S3**). Together, these two experimental systems span the physiological size range of budding yeast mother cells, which can grow to over 200 fL as they age ^30^.

**Figure 2:**
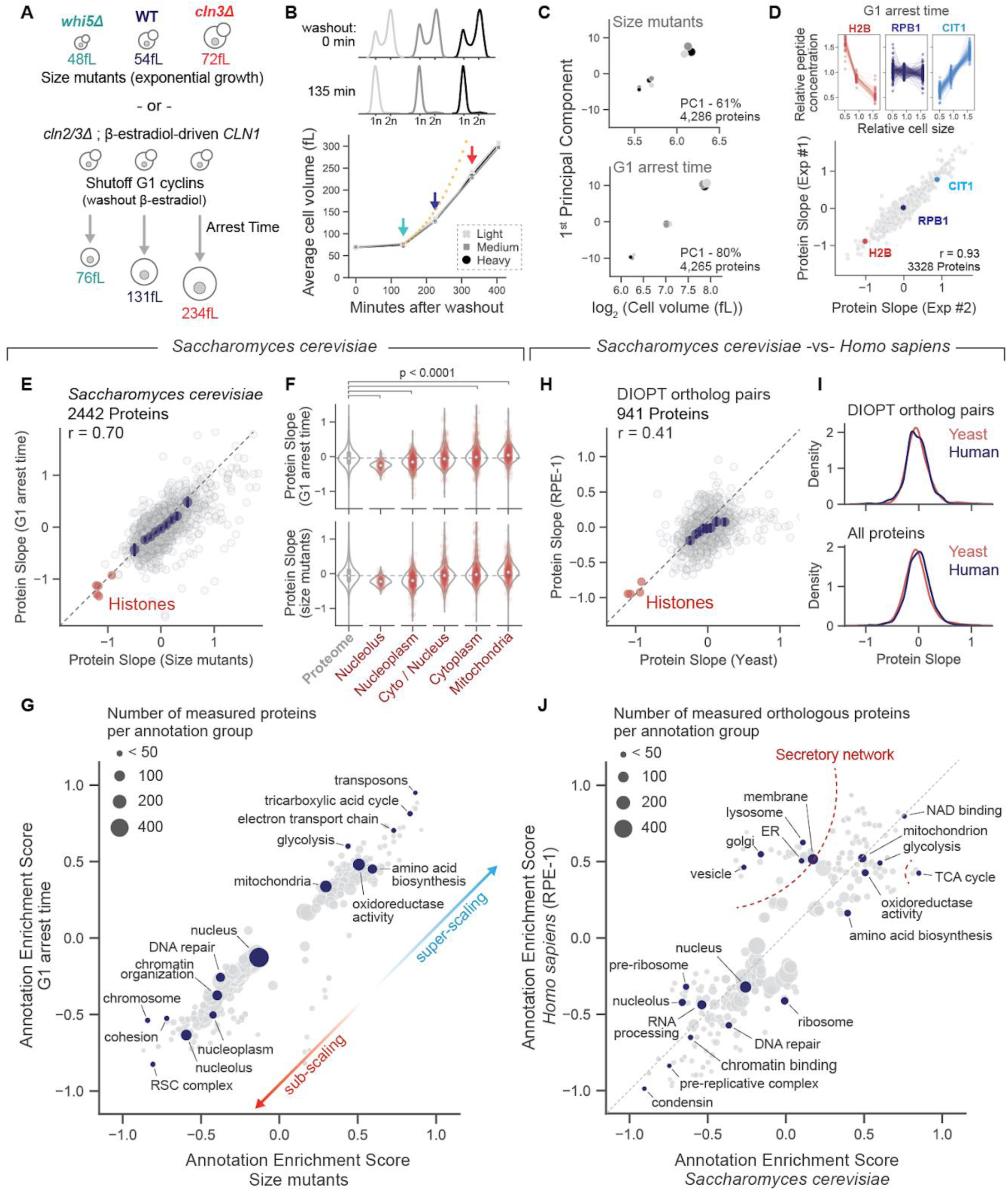
Proteome size-scaling is conserved from yeast to human cells. **(A)** Orthogonal strategies to generate populations of different-sized budding yeast. Mean cell sizes were determined using a Coulter counter (see Figure S2 for size distributions from 3 replicate experiments). **(B)** G1 arrest time course using a conditional G1 cyclin expression system. Cells arrest in G1 after removal of β-estradiol from the culture medium. Growth during arrest results in increasing cell size. Colored arrows denote the time points measured by proteomics. Grayscale colors represent the SILAC label. Light, medium, and heavy SILAC labels were swapped for three replicate experiments. **(C)** Principal component analysis of the proteome measurements of small-, medium-, and large-sized cells. The 1st principal component is plotted against mean cell volume. Dot size represents mean cell volume. Colors represent the SILAC label. SILAC labeling orientation for small, medium, and large cells was swapped for replicate experiments. All three replicate experiments are plotted together. **(D)** Top: SILAC channel intensities are plotted for three proteins that exemplify different size scaling behaviors. Each dotted line represents an independent peptide measurement. Bottom: Protein slope values from two replicate experiments. Only proteins with at least 4 independent peptide measurements in both experiments are shown. r value denotes Pearson correlation coefficient. **(E)** Correlation of protein slope values derived from two orthogonal experimental systems. Each dot represents the average slope value measured from 3 replicate experiments with each size separation strategy. Only proteins with at least 12 independent measurements (see methods) in both set of 3x replicates are shown (see Table S3 for all protein measurements). Core histone proteins are shown in red. The identity line is dashed. Blue dots are x-binned data and error bars represent the 99% confidence interval. r value denotes Pearson correlation coefficient. **(F)** Size-scaling behavior of individual proteins strictly parsed by subcellular localization (see methods). **(G)** 2D annotation enrichment analysis using the protein slope values calculated from the two orthogonal experimental systems. Each dot is an annotation group, and the position of the dot is determined by the mean slope value of the proteins in the group (rank-based). Positive and negative enrichment scores indicate groups of super-scaling and sub-scaling proteins, respectively. Table S4 contains a complete list of enrichment scores for significantly super- or sub-scaling GO terms. **(H)** Correlation of protein slope values for budding yeast (G1 arrest time) and cultured human cells (data from Lanz *et al.*^19^). Proteins are paired by sequence orthology using DIOPT (see methods). Core histone proteins are shown in red. The identity line is dashed. Only proteins with a minimum of 7 independent measurements in both the budding yeast and human datasets are considered. Blue dots are cohorted x-binned data and error bars represent the 99% confidence interval. r value denotes Pearson correlation coefficient. **(I)** Distribution of Protein Slope values derived from budding yeast and cultured human cells. The top distribution only considers the proteins paired by orthology and plotted in Figure 2H. The bottom distribution considers all measured yeast and human proteins. Only proteins with a minimum of 7 independent measurements in both the budding yeast and human dataset are considered. **(J)** 2D annotation enrichment analysis using the Protein Slope values paired by sequence ontology between budding yeast and cultured human cells. The identity line is dashed. Each dot is an annotation group, and the position of the dot is determined by the mean slope value of the proteins in the group (rank-based). Positive and negative enrichment scores indicate groups of super-scaling and sub-scaling proteins, respectively. Table S4 contains a complete list of enrichment scores for significantly super- or sub-scaling GO terms.

By manipulating yeast cell size in two different ways, we found the proteome remodeled continuously with changes in cell size (**Figure 2C**). To quantify these changes, we calculated slope values to describe how the relative concentration of each measured protein changes as a function of cell size (**Figure 1B, 2D and S3E**). Size-dependent proteome changes were similar in both experimental systems (**Figure 2E and S3G; Table S3**), indicating that the proteome is continuously remodeled throughout a ∼5-fold size range in budding yeast (from *whi5Δ* to large G1-arrested cells). The concentrations of *most* proteins measurably changed with size (**Figure S3H**). The absolute abundance of a protein did not predict its concentration changed with cell size (**Figure S3F**). Like in mammalian cells, the proteome content of different organelles scaled differently as cell size increased (**Figure 2F**), with the nuclear/nucleolar proteins becoming more dilute and the mitochondrial proteins becoming more concentrated. 2D annotation enrichment^31^ revealed a range of scaling behaviors across many different annotation groups (**Figure 2G; Table S4**). Our proteome measurements were in general agreement with what was previously measured in excessively large, arrested cells ^32^. Importantly, we also confirmed that size-dependent proteome changes were not the result of auxotrophy in our SILAC-compatible yeast strains or the depletion of a limiting nutrient in the media (**Figure S4**).

When grouped by subcellular localization, the size-dependent proteome changes in budding yeast were broadly similar to those of mammalian cells (**Figure 1I and 2F**). To quantitatively compare size scaling in yeast and mammalian cells, we matched individual yeast and human proteins based on sequence orthology (DIOPT) ^27^ and compared how their concentrations changed with cell size. The regression of protein slope values from orthology-matched yeast and human proteins produced a strongly significant correlation (p < 0.0001) (**Figure 2H; Table S2**). The degree to which the proteome changes with cell size is also the same in yeast and humans (**Figure 2I**). Moreover, a 2D annotation analysis of the orthology-matched protein slopes revealed broadly similar size-scaling behaviors for most protein annotation groups. The most prominent difference between how the proteome scales with size in yeast and mammalian cells was in the proteins in the secretory network, which super-scaled exclusively in mammalian cells (**Figure 2J; Table S4**). Nevertheless, we find the degree of conservation of proteome size-scaling in yeast and human cells more striking than the differences.

### Transcriptional and post-transcriptional mechanisms underlie the size-scaling behavior of individual budding yeast proteins

After identifying widespread size-dependent proteome changes in budding yeast, we next sought to investigate the underlying molecular mechanisms. We first leveraged existing bioinformatic data to determine whether size-dependent proteome changes are regulated transcriptionally. Because budding yeast mutant strains have a wide range of average cell sizes, we re-examined the microarray data from 1,484 yeast mutant strains as analyzed previously ^7^. For each transcript, we regressed its relative concentration in each mutant strain against the mean cell size of that strain to calculate a Pearson r correlation value (**Figure 3A**). These transcript-vs-size Pearson r values should reflect the size-scaling behavior of individual mRNA transcripts. Indeed, transcript-vs-size Pearson r values strongly correlated with our proteomic measurements (**Figure 3B and S5A**), indicating that size-dependent proteome changes are regulated transcriptionally. Importantly, the size differential across the 1,484 strains used in our analysis is the result of mutations to a diverse set of cellular pathways. Thus, although our proteomic measurements primarily relied on manipulating the G1/S transition (**Figure 2A**), the proteome changes we observed are a general property of cell size rather than an idiosyncratic effect of alterations to the cell cycle. Consistent with this observation, cell cycle-regulated genes were not enriched as either sub- or super-scaling proteins (**Figure S5B**).

**Figure 3:**
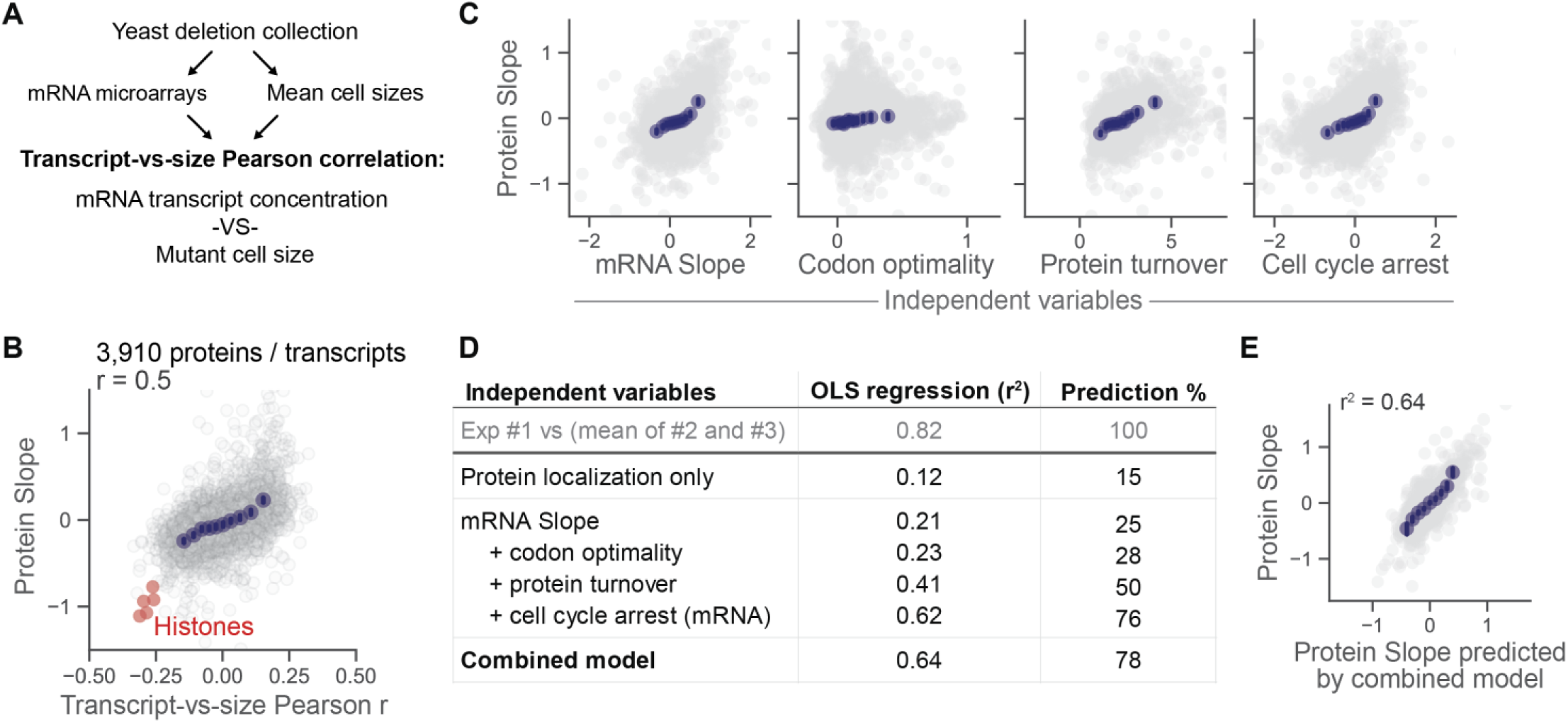
Transcriptional and post-transcriptional mechanisms underlie the size-scaling behavior of individual proteins. **(A)** Strategy to determine the size scaling behavior of individual transcripts from the mRNA microarray measurement of 1,484 yeast mutant strains. For each measured transcript, its relative concentration in each mutant strain was regressed against the size of that mutant strain. Pearson r values were calculated that reflect the size-scaling behavior of individual mRNA transcripts, as was done previously ^7^. **(B)** Correlation of size-dependent proteome changes (G1 arrest time) with the Pearson r values calculated in (A). Histone proteins and transcripts are highlighted in red. Blue dots are cohorted x-binned data and error bars represent the 99% confidence interval. r value denotes Pearson correlation for the plotted data. **(C)** Correlation of protein slope with multiple model variables. mRNA slope was calculated from transcript TPM values for G1 arrested cells. “Cell cycle arrest” represents the transcript change from initial cell cycle arrest while “mRNA slope” represents the size-scaling of the transcript throughout the course of the G1 arrest. Codon score and protein turnover values were taken from YeastMine ^35^ and a previous study ^34^, respectively. **(D)** Linear regression models to predict protein slope values based on the variables in (C). The protein localization model is based on binary association with a select set of subcellular compartments (like in Figure 2F). Codon score, protein turnover, and cell cycle arrest data were iteratively incorporated to a model derived from mRNA slopes. The combined model represents all the independent variables in (C) as well as subcellular localization. Correlation between biological replicates was used as a benchmark for the maximum predictive power. **(E)** Comparison of protein slopes predicted from the combined linear model and protein slopes measured from G1 arrested cells (Figure 2).

To determine the degree to which size-dependent proteome changes are regulated transcriptionally, we performed RNAseq analysis on different sized cells and calculated mRNA slope values analogous to the protein slopes described in Figure 1B. mRNA slopes strongly correlated with corresponding protein slopes indicating that much of the protein level size-scaling is due to changes in gene expression (**Figure 3B**). This finding is consistent with the few cases of differential scaling studied in-depth, Whi5 and histones, where protein sub-scaling was due to mRNA sub-scaling ^6,7,33^.

In addition to transcript concentration, we investigated whether post-transcriptional variables contributed to proteome size scaling using linear models. We found that a protein’s turnover ^34^ and codon affinity ^35^ both significantly correlated with the Protein Slope (**Figure 3B**). Iterative incorporation of each parameter (mRNA slope, protein turnover, and codon affinity score) significantly improved the model with minimal collinearity (**Figure 3C and S5C**). Using the correlation between biological replicates as a benchmark for the maximum size-dependent variation any model could predict, we find that our composite model predicts ∼80% of this maximum size-dependent variance (**Figure 3D**). Taken together, our linear models indicate that size-dependent proteome remodeling is driven both pre- and post-transcriptionally.

### Size-dependent proteome remodeling is mostly independent of metabolic state

As budding yeast grow larger, many proteins associated with respiratory metabolism become increasingly concentrated. For example, proteins involved in the tricarboxylic acid cycle and electron transport chain are among the most super-scaling in our dataset (**Figure 2C**). Since the upregulation of respiratory metabolism typically happens when yeast are starved of glucose (*e.g.*, the diauxic shift), we wondered whether the size-dependent proteome changes could be explained by a starvation-like shift to respiratory metabolism that occurs with cell enlargement despite the presence of excess glucose in the media.

To test whether such a shift in metabolic state could explain size-dependent proteome remodeling, we first measured the proteome of yeast growing on a non-fermentable carbon source and compared it to the proteome of yeast growing on glucose (**Figure 4A and S6A,B**). We calculated a concentration ratio (respiration:fermentation) for 4,162 yeast proteins (**Figure 4A and S6D; Table S5**). As expected, the fraction of the proteome dedicated to respiratory metabolism was significantly higher in yeast utilizing ethanol/glycerol, the non-fermentable carbon source (**Figure 4B**). Interestingly, nearly half of the proteins measured in our analysis changed in concentration by 2-fold or more between carbon sources. The propensity of high copy number proteins (1st abundance quartile) to change in concentration was the same as the low copy number proteins (4th abundance quartile) (**Figure 4C and S6C**). In other words, the proteome is extensively remodeled when shifting between these two carbon sources. Yet despite the large difference in baseline proteome content between fermenting and respiring yeast, the way in which the proteome remodeled with cell size within these two metabolic states was remarkably similar (**Figure 4D and S6E,F; Table S3**). Interestingly, some of the protein groups that were most upregulated in respiring yeast (**Figure 4B**) were no longer super-scaling with cell size when the yeast were grown in ethanol/glycerol media (**Figure 4E,F; Table S4**). This may suggest that the baseline upregulation of these protein groups in yeast growing in ethanol/glycerol is distinct from the super-scaling behavior present in glucose. Overall, these results show that size-dependent proteome remodeling is mostly independent of a cell’s metabolic state, though the size-scaling behavior of certain protein groups may depend on growth conditions.

**Figure 4:**
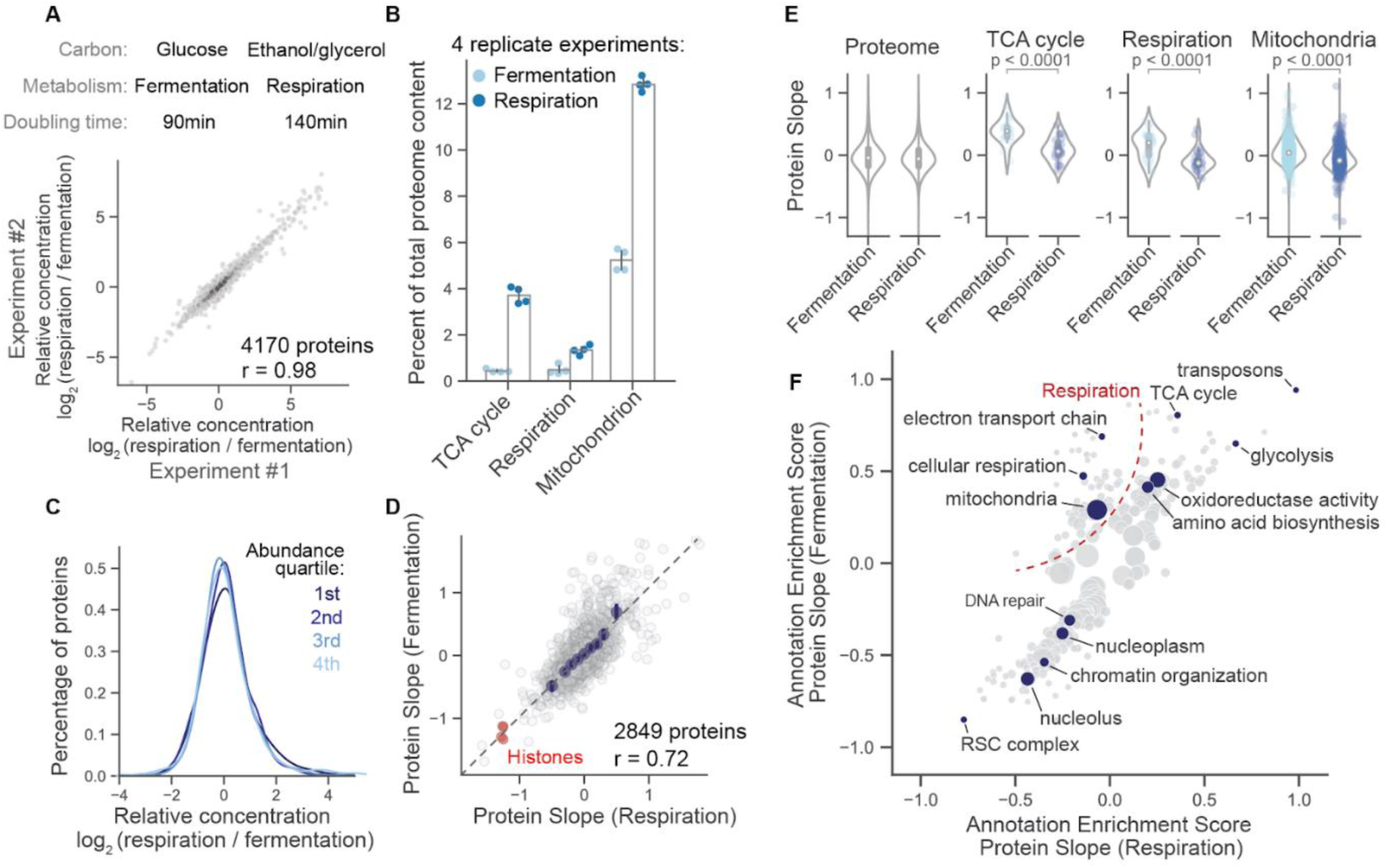
Metabolic state has a large effect on proteome composition, but only a small effect on how the proteome changes with cell size. **(A)** Proteome comparison of yeast growing asynchronously in media containing glucose or ethanol/glycerol as a carbon source. Relative concentration ratios (respiration / fermentation) for individual proteins measured in two replication experiments are plotted (see Figure S6 for supporting information). R value denotes the Pearson correlation coefficient. Data density is shaded. **(B)** Fraction of the proteome dedicated to the indicated GO term. Proteome fraction is calculated from the summed intensity of all the peptides from all proteins possessing the indicated GO term divided by the total peptide ion intensity. Each dot represents an independent biological replicate. **(C)** Distribution of relative concentration changes (Respiration / Fermentation) for proteins with different expression levels. Protein abundance was separated into quartiles using summed intensity for all peptide measurements for a given protein. The 1st and 4th quartiles represent the 25% most and least abundant proteins in the dataset, respectively. X-axis represents the mean ratio from the experiments in (A). **(D)** Correlation protein slope values derived from yeast growing in ethanol/glycerol- and glucose-containing media. Core histone proteins are shown in red. Glucose-derived protein slopes are from the G1 arrest time experiment in Figure 2. The same experimental system was used to generate the ethanol/glycerol-derived Protein Slope values (see Figure S5). Only proteins with a minimum of 4 independent measurements in each dataset are considered. Blue dots are x-binned data and error bars represent the 99% confidence interval. r value denotes the Pearson correlation coefficient. **(E)** Violin plots depicting the mean and distribution of protein slope values for the indicated annotation groups. p-values comparing the distributions are shown above each comparison. **(F)** 2D annotation enrichment analysis (see methods) using the protein slope values derived from glucose and ethanol metabolizing yeast. Each dot is an annotation group, and the position of the dot is determined by the mean slope value of the proteins in the group (rank-based). Positive and negative enrichment scores indicate groups of super-scaling and sub-scaling proteins, respectively. Table S4 contains a complete list of enrichment scores for significantly super- or sub-scaling GO terms.

### The general stress response super-scales with cell size in budding yeast

Having found that size-dependent proteome remodeling is mostly independent of the external nutrient environment, we wondered whether it might instead be triggered by an intrinsic stress associated with cell enlargement. To investigate this possibility, we compared our size-dependent proteome changes to those observed in response to several different stress conditions (**Figure 5A**). This analysis revealed substantial overlap between size-dependent proteome remodeling and several acute stresses, including osmotic stress ^36^, starvation ^37^, and mTOR inhibition (this study) as well as a previously reported signature of associated with slow steady-state growth (**Figure 5A**) ^38,39^. In addition to their correlation with size-dependent proteome changes, all these -omics datasets show strong cross-correlation (**Figure 5A**): A shared pattern of gene expression changes known the environmental stress response (ESR). Thus, large cell size is associated with potent ESR activation.

**Figure 5:**
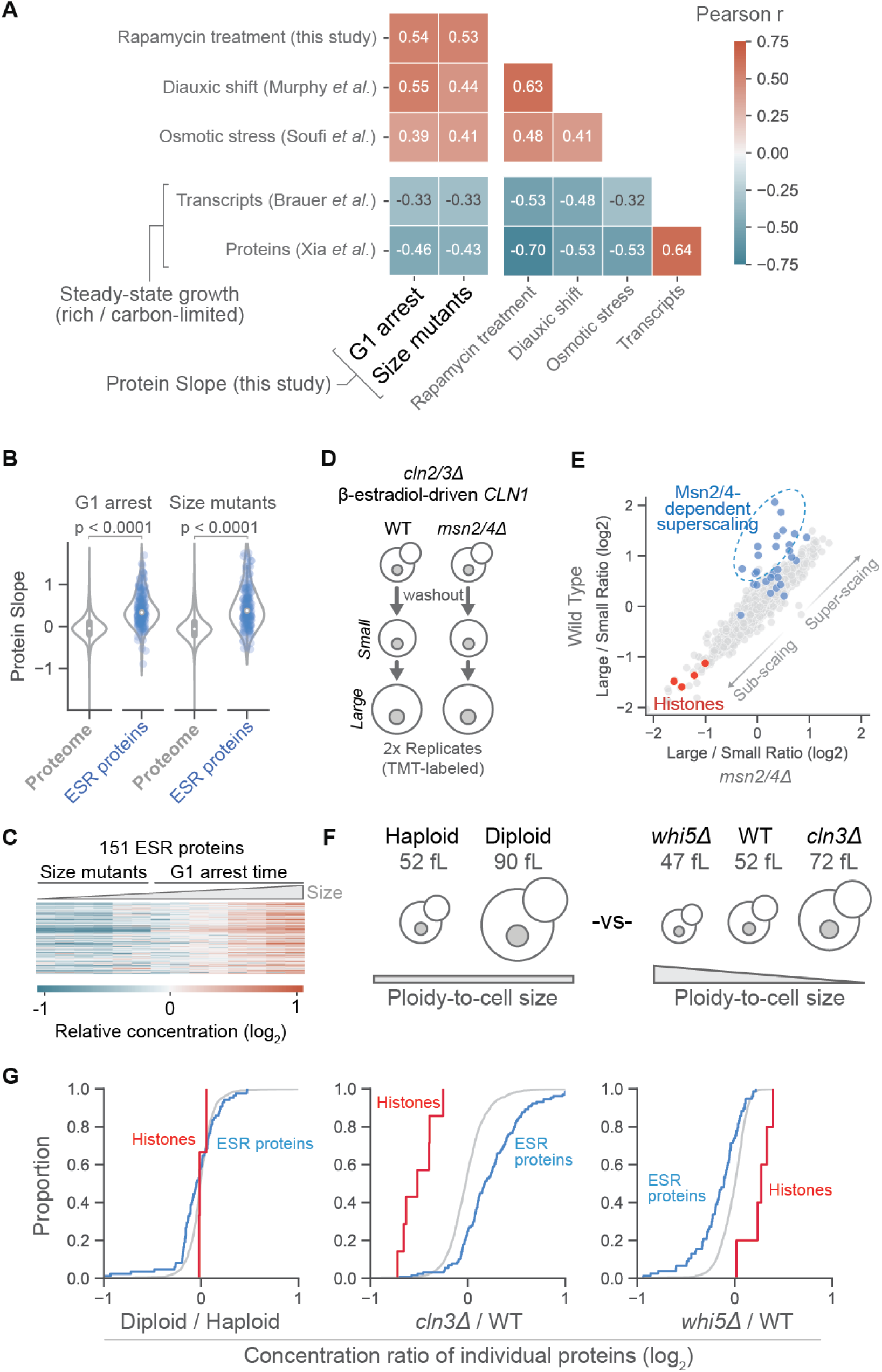
The general stress response super-scales with cell size. **(A)** Matrix showing the cross correlation of size-dependent proteome changes and the proteome/transcriptome changes that occur in response to different stresses. For the Murphy *et al.* dataset, protein concentration ratios were calculated from cultures in either the log or lag phases of the diauxic shift in budding yeast. For Soufi et al, concentration ratios were calculated before and after 20-minute exposure to 400mM NaCl. For the rapamycin dataset, concentration ratios were calculated before and after a 150-minute treatment of 1µM rapamycin. The two “steady state” datasets used chemostats to compare the transcriptome and proteome in glucose replete and limited growth conditions. We used the “Growth Rate Slope ’’ reported by Brauer *et al.* and equivalent metric from Xia et al. for the transcript- and protein-level correlations, respectively. The steady-state datasets are anti-correlated with the protein slope because the “Growth Rate Slope ’’ reported by Brauer *et al.* reflects (no stress / stress) ratio orientation, while the datasets with acute stress induction have a (stress / no stress) orientation. **(B)** Violin plots showing protein slope values for the genes that are upregulated during the environmental stress response (ESR) ^38^ indicate that this gene set is predominantly super-scaling. **(C)** Heatmap depicting the size-dependent increase in the concentration of ESR proteins across a ∼4-fold range of budding yeast sizes (Table S6). Different-sized cells from both experimental systems (G1 arrest time and size mutants) were labeled with TMT and quantified together. Mean cell size is increasing from left to right (*whi5Δ* to the final time point of the G1 arrest). Two replicates for each mean size are shown as paired columns. **(D)** G1 arrest time course experiments were performed in mutant strains that lack the ESR transcription factors Msn2 and Msn4. **(E)** Correlation of size-dependent proteome changes in cells with and without Msn2 and Msn4. ESR proteins whose baseline expression in small WT cells is dependent on Msn2/4 (Figure S6D) are highlighted in blue. Histones are highlighted in red. **(F)** A strategy to generate yeast of different sizes that have a similar cell size-to-ploidy ratio. **(G)** Mean concentration ratio from a label-swapped SILAC comparison of the indicated strains (Table S7). Gray cumulative distribution represents all measured proteins. ESR-associated and histone proteins are highlighted in blue and red, respectively.

Is large cell size a general stress? In addition to our observations with G1-arrested cells (**Figure 5B**), other recent work also observed ESR engagement in large, arrested cells ^32,40^. Interestingly, the upregulation of the ESR was not dependent on cell cycle arrest, since the ESR proteins super-scaled to the same degree in asynchronously proliferating size mutants as they did in G1 arrested cells (**Figure 5B**). Searching for potential mechanistic explanations, we found that the degree of ESR activation increased in near linear proportion to cell volume across both the asynchronously growing size mutants and G1 arrested cells (**Figure 5C and S7A; Table S6**). Mutating Msn2 and Msn4, two primary facilitators of the ESR ^41^, attenuated the super-scaling of some but not most ESR-associated proteins (**Figure 5C and S7B,C,D; Table S6**). ESR activation did not occur when increased cell size was accompanied by increased ploidy (**Figure 5G and S7E; Table S7**), suggesting that this response is triggered by changes in the ploidy-to-cell size ratio. Interestingly, the deletion of *WHI5*, which reduces cell size but not growth rate, decreased the expression of ESR-associated proteins relative to WT cells (**Figure 5G and S7E; Table S7**).

### Age-associated proteome changes are largely explained by the age-associated increase in cell size

The induction of a stress response upon cell enlargement may have important implications for yeast replicative aging. Over the course of their lifespans, budding yeast mothers produce ∼25 daughters, growing progressively larger during each G1 phase before they permanently exit the cell cycle ^42^. Given our findings above, we hypothesized that a substantial portion of the age-associated proteome changes could be explained by increased cell size.

To test this hypothesis, we measured how the proteome changes with age and compared these data to the proteomic signature of increased cell size. To do this, we combined a biotin-streptavidin affinity purification procedure with a genetic selection scheme known as the ‘mother enrichment program’ to isolate aged yeast (**Figure 6A,B**) ^42^. This allowed us to isolate and quantify the proteomes of aged mother cells and their daughters. In concordance with prior observations, we observed substantial differences between the proteomes of old and young yeast, correlating most strongly with a recently a recently published study (**Figure S8D**) ^43^. These data correlated remarkably well with the proteomic signature of increased cell size (**Figure 6D and S8C**) despite differences in strain background. Moreover, most age-associated changes in the proteome were also present in the daughters of old cells (**Figure S7A**), which have a very similar lifespan to the daughters of young cells. These observations suggest that many age-associated proteome changes may be entirely explained by the increased size of old mother cells. These include sentinel age-associated changes, such as an apparent decrease in histone proteins (**Figure 6C and S8B**) often referred to as ‘nucleosome loss’ ^44^. Our data suggest that this phenomenon arises from the increased size of older yeast rather than an age-associated “loss” of existing histones.

**Figure 6:**
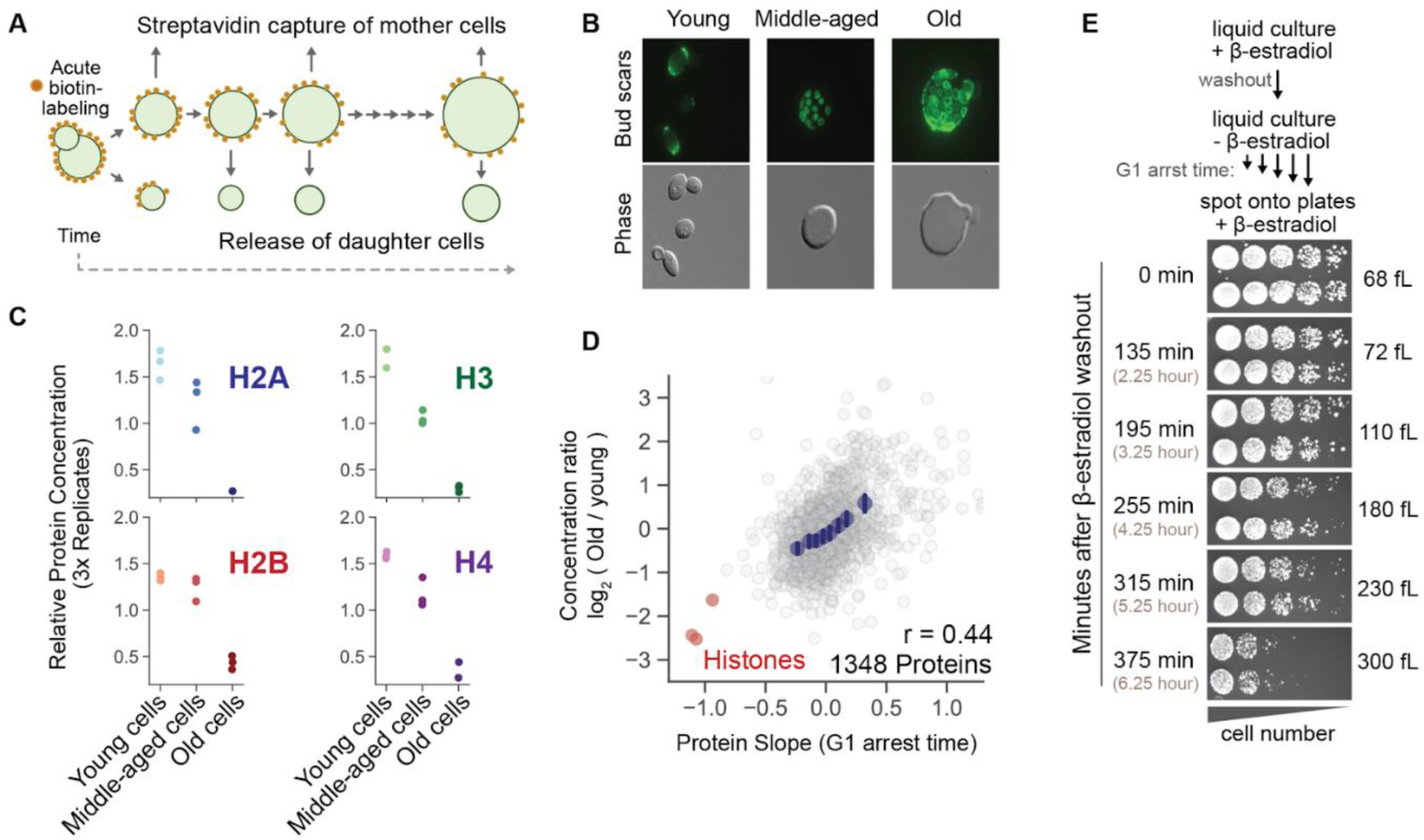
Age-associated proteome changes are largely due to the age-associated increase in cell size. **(A)** A strategy to enrich for replicatively-aged yeast based on the mother enrichment program (see methods). Cell size increases with age in both mother and daughter cells ^42^. After incubating labeled cultures for different periods of time, we measured the proteomes of young (0h), middle aged (24h), and old yeast (48h) using TMT (see methods). **(B)** Example fluorescent and phase images of young, middle-aged, and old yeast. Bud scars are fluorescently labeled (WGA-FITC). Images illustrate the increased size of aged budding yeast mothers. **(C)** Relative concentrations of core histone proteins in young, middle-aged, and old yeast. Each dot corresponds to 1 of 3 biological replicate experiments. **(D)** Correlation of age-associated and size-dependent proteome changes. Protein slope value is from the G1 arrest time experiment in Figure 2. Only proteins with a minimum of 4 independent measurements in each dataset are considered. Histone proteins are highlighted in red. Blue dots are x-binned data and error bars represent the 99% confidence interval. r value denotes Pearson correlation coefficient. **(E)** Spot assay showing the decline in proliferative capacity in very large cells. Cells are arrested and enlarged as described in Figure 2 (“G1 arrest time”). The mean cell size at the time of spotting on +β-estradiol plates is shown in fLs. Each plate was imaged after 40 hours of growth at 30°C.

If many age-associated proteome changes are a consequence of increased cell size, to what extent are the pathologies associated with aging, most notably senescence, due to large cell size? A previous study used a temperature sensitive *cdk1-ts* allele to arrest and enlarge yeast for a prolonged period of time (6 hours) ^32^. When the enlarged yeast were returned to the permissive temperature, a proliferation defect was observed ^32^. Here, we performed an analogous arrest and release experiment using our hormone-inducible system (**Figure 6E**). We found that when our G1-arrested cells surpassed a size threshold (∼200fL), their viability sharply declined (**Figure 6E**, bottom most panel). Interestingly, the size range around which a significant fraction of G1 cells permanently lost the ability to re-enter the cell cycle corresponded to the upper size range of old haploid yeast mother cells ^30^. This observation supports the hypothesis that cell size is an important determinant of budding yeast lifespan, rather than simply a correlate ^30,45^.

### Genome limitation drives starvation-like proteome remodeling in eukaryotes

Our experiments so far show that the concentration of the eukaryotic genome (*i.e.*, ploidy-to-cell size ratio) drives changes in proteome composition (**Figure 1, 2, and 5**) and that increasing cell size is associated with the induction of a stress response (**Figure 5**). Another phenotype often linked to the general stress response in yeast is a slow growth rate. Consistent with the connection between large cell size and the induction of a stress response, the specific growth rate (mass added per unit mass) of G1 yeast or human cells declines as cell size increases (**Figure 2B and S3B**) ^46–48^. Moreover, size-dependent proteome changes in both yeast and human cells strikingly resemble those that occur after acute growth inhibition by exposure of cells to the TOR inhibitor rapamycin (**Figure 7A; Table S9**) ^47^. Taken together, these results support a model where the genome becomes limiting for growth in large cells and that this limitation elicits a starvation-like proteome phenotype (**Figure 7A,B**). Consistent with this model, the growth defects and proteome remodeling associated with cell enlargement are averted when an increase in size is accompanied by an increase in ploidy (**Figure 5G and 7B**) ^26,32^.

**Figure 7:**
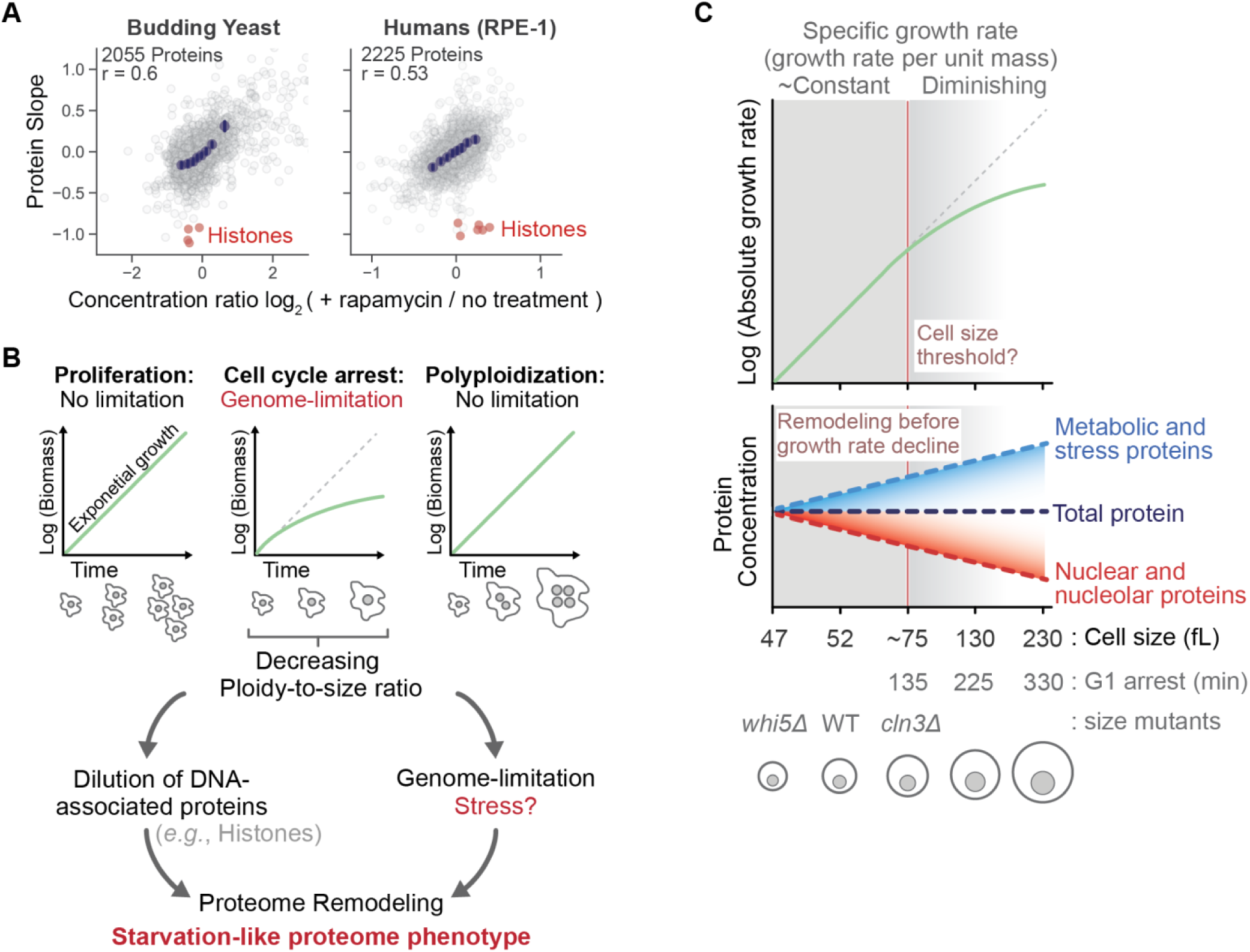
Genome concentration is a universal determinant of proteome content in growing cells. **(A)** Correlation of protein slopes values and the relative concentration changes in response to acute growth inhibition of asynchronously proliferating cells (saturating dose of rapamycin for 150 minutes and 24 hours in budding yeast and human cells, respectively). The correlation in human cells is re-plotted from published work^47^. Blue dots are x-binned data and error bars represent the 99% confidence interval. r value denotes the Pearson correlation coefficient. **(B)** Model for how increasing cell size drives proteome remodeling. Schematic illustrates how growth is genome-limited. Dotted identity lines represent exponential biomass accumulation. Decreasing genome concentration (*i.e.*, ploidy-to-cell size ratio) drives proteome remodeling in distinguishable ways. **(C)** Schematic showing how the proteome remodels continuously with increasing cell size even though the specific growth rate only declines once yeast pass a certain size threshold. X-axis is not drawn to scale.

## DISCUSSION

It was long thought that the concentrations of individual proteins remain mostly constant as cells grow larger, in part because total protein amount scales in direct proportion to cell size ^49^. Recent proteomic analyses have refuted this paradigm by identifying widespread size-dependent changes in the concentrations of individual proteins ^19,20^. Here, we identify conserved principles for size-dependent proteome remodeling by extending our analysis to evolutionarily divergent eukaryotes. We find that in mammalian and yeast cells, the proteome remodels continuously as cells grow larger. These findings are important because they provide a simple and universal rationale for the regulation of cell size. Namely, the further a cell deviates from its target size, the more biochemically different it becomes.

### Many DNA-associated proteins are maintained in proportion to DNA amount

In every experiment where we manipulated cell size in eukaryotes, the relative change in the concentration of histone proteins matched the relative change in the concentration of the genome. A mechanism for how this type of regulation is achieved has recently been proposed. It was found that transcription of the histone loci is tightly coupled to S phase in an amount independent of cell size ^7,33^. In addition, histones produced in excess because of delays in DNA replication are post-translationally degraded ^50^, resulting in protein amounts that are always proportional to the amount of DNA. In addition to the histone proteins, the concentrations of several other DNA-associated proteins (*e.g.*, cohesion, condensin, and chromatin remodeling complexes) also aligned with changes in genome concentration (**Figure 2F**), though it is unclear if they are subject to the same mechanisms of regulation as histones. The maintenance of a strict stoichiometry between the genome and tightly DNA-bound proteins, such as histones, may prevent an excess of these proteins in the chromatin. Notably, the overexpression of histone proteins is often toxic ^51^.

### Is cell enlargement *per se* stressful?

Perhaps the most striking set of super-scaling proteins in budding yeast are those associated with the general stress response (also referred to as the environmental stress response or ESR) (**Figure 5B**). The notion that cell enlargement is intrinsically stressful is consistent with connections between excessive cell enlargement and senescence in cultured mammalian cells. Senescence is the state of irreversible (or durable) cell cycle arrest in which cells fail to divide in response to mitogens or oncogenic stimuli ^52^. Mounting evidence suggests that cell enlargement promotes senescence-associated proteome changes and promotes cell cycle exit ^4,19,20,32,53–55^, but the mechanism by which this happens is unclear. Like in yeast, excessive cell size could trigger a stress response leading to increased expression of the Cdk inhibitors p16 or p21, which are associated with senescence ^52^. Indeed, multiple independent groups have now reported that p21 is upregulated with cell enlargement in various human cell lines ^19,20,56–59^. One potential source of p21 induction is the increased susceptibility to DNA damage in large cells ^19,56^, which may, in part, result from the replication stress caused by the sub-scaling of DNA replication and repair proteins ^60^ or an inability to initiate DNA repair ^56^. Other proposed sources of stress during cell enlargement include cytoplasmic dilution ^32^ and osmotic stress ^58^. Consistent with links between cell enlargement and senescence identified *in vitro*, recent work has shown that the proliferative potential of only slightly enlarged hematopoietic stem cells is severely compromised *in vivo* ^61^.

In budding yeast, ESR induction is tightly linked to decreases in the cellular growth rate ^38^. Surprisingly, we find size-dependent changes in ESR expression even in cases where the cellular growth rate is not changing. For example, the ESR is upregulated in an asynchronously proliferating *cln3Δ* strain (**Figure 5G**), which has a doubling time close to wild type (**Figure S3A**). Similarly, we observed a reduction of ESR expression in the smaller *whi5Δ* strains (**Figure 5G**), which also proliferate normally (**Figure S3A**). ESR induction is likely caused by increased cell size rather than by cell cycle arrest since the degree of ESR induction is directly proportional to differences in size regardless of whether cells were arrested in the cell cycle (**Figure 5B**). It is possible that even wild type yeast experiences some basal-level stress due to their size, since the concentration of ESR-associated proteins can be decreased through mutations such as *whi5Δ* that reduce cell size but do not affect proliferation rates (**Figure S3A**). Thus, it is possible that the super-scaling of stress response proteins may not be directly driven by stress. Rather, the super-scaling of the ESR could represent an evolved mechanism to passively prepare smaller daughters for the stress that will come as they become progressively larger with age. Regardless of its causality, a recent study demonstrated that ESR induction in large, arrested yeast cells reduces the decline in their viability ^40^. This implies that the ESR response in progressively larger cells is beneficial.

### Genome dilution elicits a starvation-like proteome phenotype

The induction of general stress response has also been attributed to changes in the cellular growth rate in budding yeast. Attenuating cell growth by limiting the availability of any single nutrient results in the upregulation of ESR transcripts ^38^. Moreover, the proteome remodeling that occurs in response to acute stress is similar to how the yeast proteome changes when cells are starved or grown at steady-state in carbon-limiting media (**Figure 5A**). It is unclear how the general stress response is coupled to changes in growth rate, but this coupling is functional in certain scenarios since slow growing cells are more resistant to certain types of acute stress ^62^.

One possibility is that the proteome changes we observe with increasing cell size are connected to the declining specific growth rate (growth rate per unit mass) in large cells (**Figure 2B and S3B**) ^46^, as was recently proposed for human cells ^47^. Consistent with this idea, we find that the dilution of the yeast genome elicits a starvation-like proteome phenotype (protein slope vs diauxic shift, **Figure 5A**), and, in both yeast and mammals, size-dependent proteome changes are strikingly similar to those that occur after cell growth is acutely inhibited (**Figure 7A**) ^47^. The observation that increasing cell size proteomes a starvation-like proteome phenotype suggests that cell growth is intrinsically limited by the genome in a manner analogous to a limiting nutrient (**Figure 7B**). The starvation-like proteome changes in large cells may also explain the allometric size-scaling of mitochondrial membrane potential, which was described previously ^63,64^.

The current models for how different proteome “sectors” are reallocated when cells are grown in different nutrient conditions typically describe a linear relationship between proteome content and the cellular growth rate. For example, the proteome sector allocated to protein synthesis decreases (as a fraction of the total proteome) when cells are grown on poorer carbon sources in both yeast and bacteria ^39,65,66^. However, the relationship between proteome content and growth dynamics is likely more complex than the simple linear relationships described for ribosomes ^67^. For example, the widespread proteome changes that occur with increasing cell size in budding yeast precede a measurable decline in their specific growth rate (**Figure 7G**).

### The oocyte: an extreme case of genome dilution?

The largest mammalian cell, the oocyte, seemingly contradicts the principles we highlight here. The mature oocyte is exceptionally large and contains only a single copy of the genome. While cell enlargement leads to decreased proliferation and senescence in somatic cells, the oocyte obviously retains its proliferative potential. So, how does such an enormous cell defy the stress- and senescence-related defects that are observed in enlarged somatic cells? The architecture of the drosophila germline cyst ^68^ provides a clue for how the oocyte might bypass the deleterious effects of large cell size. Namely, oocyte maturation is driven by surrounding nurse cells, which directly feed the expansion of the oocyte’s cytoplasm. Each nurse cell possesses thousands of copies of the drosophila genome. Thus, the oocyte may simply rely on the genome content of “feeding” cells to grow to its final size while bypassing the defects associated with genome dilution in somatic cells.

### A model for how increasing cell size remodels the proteome

Here, we established how genome dilution impacts the proteome in yeast and mammalian cells. We hypothesize that changes in cell size-to-ploidy ratio drive proteome remodeling in distinguishable ways (**Figure 7F**). First, the amount of many DNA-associated proteins, such as histones and chromatin remodeling complexes, are maintained in proportion to the DNA amount and are therefore diluted as cells grow larger. Second, increasing cell size is associated with a general stress response. Third, dilution of the genome appears to limit cell growth in a manner analogous to a limiting nutrient, since large cell size elicits a starvation-like proteome phenotype. We anticipate future work will identify the underlying molecular mechanisms and give insight into how and why the cell’s composition remodels with its size.

## Acknowledgements

We thank Kurt Schmoller, members of the Skotheim and Elias labs for feedback on the manuscript and helpful discussions. This work was supported by the NIH through R35 GM134858 (JMS) and the Chan Zuckerberg Biohub San Francisco (JMS Investigator Award; MCL collaborative postdoctoral fellowship).

## Contributions

**MCL** designed and carried out yeast-related experiments, except for Figure 5D (performed by **LH**) and Figure 6 (performed by **DFJ** and **IZ**). **MCL** prepared samples for mass spectrometry analysis. **MCL** acquired mass spectrometry data. **MCL** and **FM** maintained the performance of the mass spectrometers. **MCL** performed all data analyses. **SZ** derived the hepatocyte primary cells and FACS-isolated the cells populations measured in Figure 1. **MPS** performed mRNA sequencing for Figure 3 and constructed the SILAC yeast strains. **MCL** and **JMS** wrote the paper. **JMS** and **JEE** supervised the study.

## Supplemental Figures and Legends

**Figure S1 - Supplement for Figure 1.**
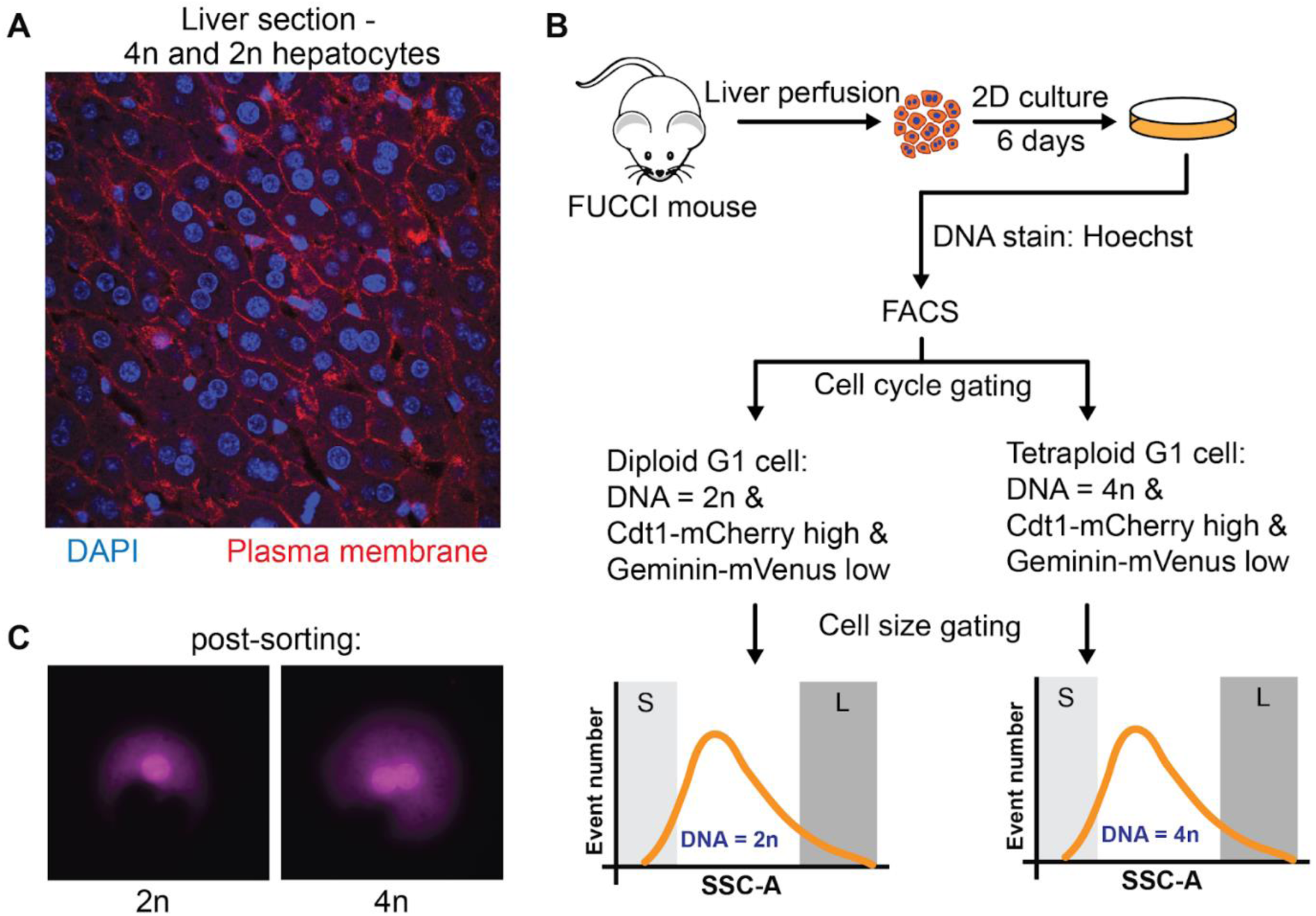
**(A)** Stained liver section illustrating the heterogeneity of hepatocyte ploidy. Mononucleated cells are diploid and binucleated cells are tetraploid. See methods for detailed description of primary cell isolation and culturing. **(B)** FACS scheme to isolate G1 phase primary hepatocyte cells of different size and ploidy. A fluorescent FUCCI cell cycle reporter was used to identify G1 phase cells. After gating for G1 cells, 2n and 4n G1 cells were differentiated using a DNA stain. Diploid and tetraploid G1 cells of different sizes were separated by the side scatter parameter. **(C)** Images of post-sorted primary hepatocytes stained with Hoechst. The mononucleated cell is diploid and the binucleated cell is tetraploid. Sorted populations were lysed and peptides were quantified using MS3-TMT proteomics (Figure 1; methods).

**Figure S2 - Supplement for Figure 2.**
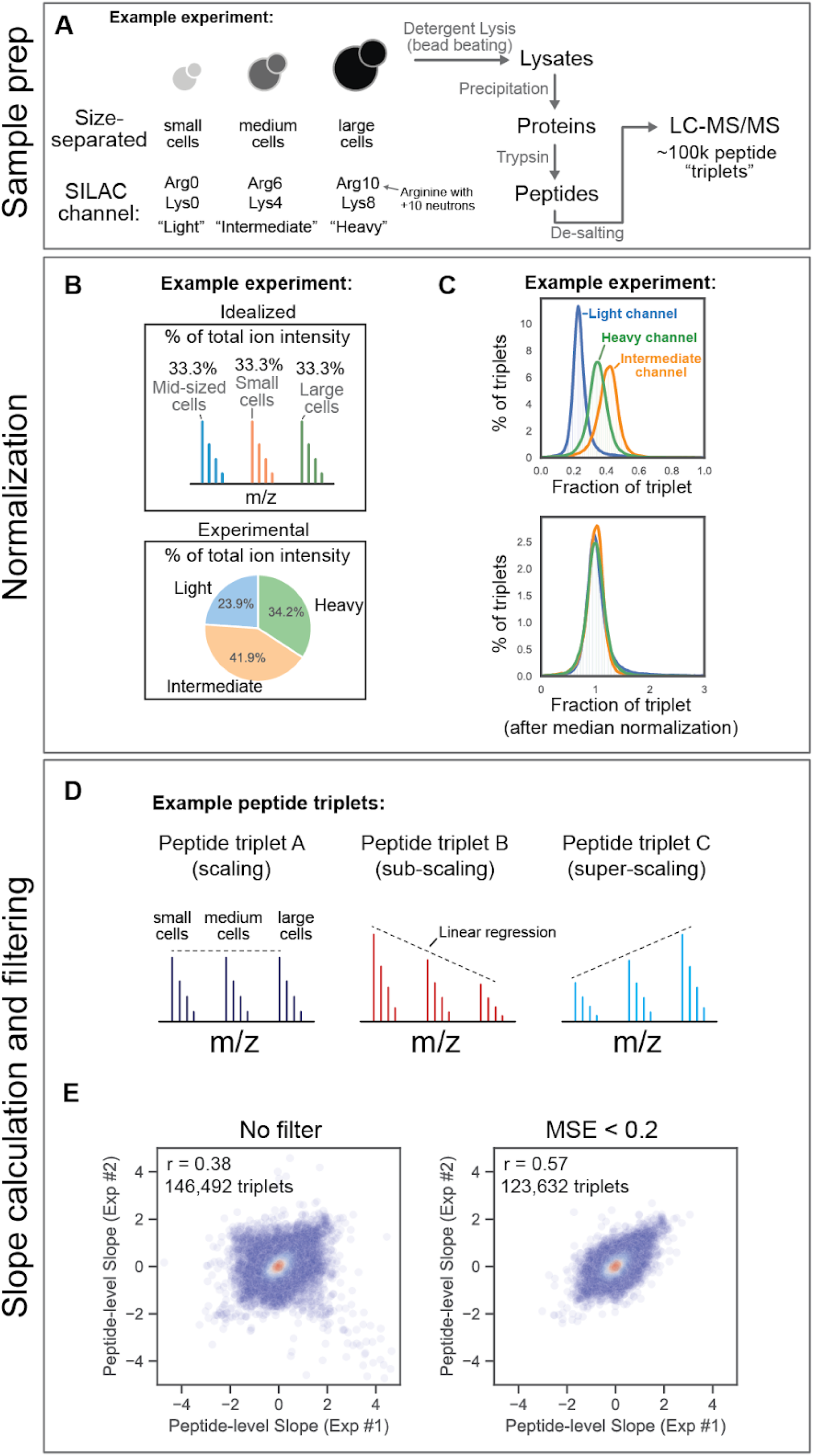
**(A)** Budding yeast of different sizes were metabolically labeled in cell culture and subjected to proteomic analysis. Strategies to separate cells by size are described in Figure 2 and S3. See methods for processing steps prior to mass spec data acquisition. **(B)** Differences in the amount of total protein contributed from the small-, medium-, or large-cell size populations were normalized as described in the methods. Rather than normalize L/H and L/M SILAC ratios separately, we normalize all three channels together so that the values in our dataset represent relative changes across all cell sizes. Data from a hypothetical example experiment is depicted. **(C)** For each individual peptide triplet, we determined the fraction of the triplet’s total ion intensity present in each SILAC channel. The distributions of these fractions were then adjusted by the median (see methods for a complete description of the normalization process). Data from a hypothetical example experiment is depicted. **(D)** Peptide slope values are calculated from a linear regression of the relative ion intensity in each SILAC channel and mean cell size. Mean cell size was determined by Coulter counter prior to mixing and lysis. The mean squared error was used to track the linear fit of each peptide triplet regression. **(E)** Correlation of peptide slopes calculated from replicate experiments before and after applying a filter for mean squared error (MSE). 146,492 unique peptide measurements were identified in two replicate experiments (“G1 arrest time” from Figure 2). A unique peptide measurement is defined by the peptide sequence, modification state, charge state, and fraction number. Loosely filtering for peptide triplets by MSE increased the correlation between biological replicate experiments.

**Figure S3 - Supplement for Figure 2.**
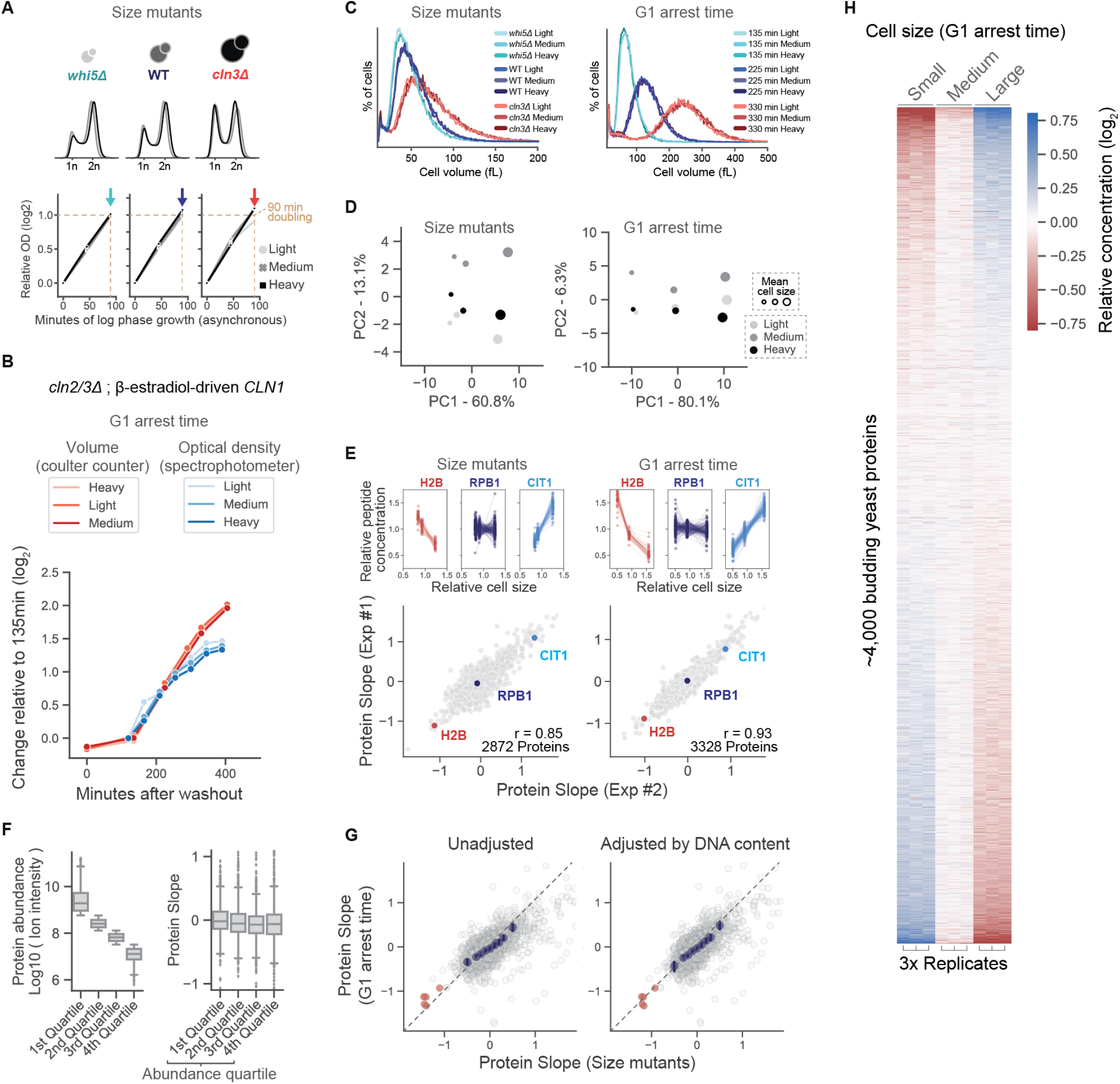
**(A)** Cell cycle distribution and steady-state growth rate is unaffected by the different SILAC labels. Cell size mutants have doubling times similar to wild type when growing asynchronously in synthetic complete media with 2% glucose (∼90min). **(B)** Growth behavior before and after G1 arrest is unaffected by the different SILAC labels. Relative change in culture density of mean cell volume is plotted in log_2_ space. The curved trajectory indicates a declining specific growth rate ∼200 minutes after washout. Measurements of volume and optical density were collected in parallel on the same cultures. Y-axis is relative to 135 min, which is when ∼95% of cells are in G1 phase. **(C)** The attainment of differentially sized cells was confirmed using a Coulter counter. Color gradients correspond to replicate cultures grown with light, medium, and heavy SILAC labels. **(D)** PCA1 vs PCA2 plotted for both orthogonal experimental systems. PC1 changes with cell size. PC2 corresponds to the SILAC labeling channel and likely reflects differences in the landscapes of analytical interference between light-, medium-, and heavy-labeled peptides. **(E)** Illustration of the protein slope calculation from relative changes in peptide concentration and cell size for both orthogonal experimental systems. See methods for a detailed explanation of the protein slope calculation. **(F)** To determine if protein abundance predicts size-scaling behavior, the proteome was first partitioned into abundance quartiles. To estimate a protein’s relative abundance, the ion intensity for all its identified peptides was summed together. We only considered the ion intensity from the “small” cell SILAC channel in each of three G1 arrest time replicate experiments. The proteome was then grouped into abundance quartiles (depicted in the left plot, see methods). The right plot depicts the distribution of slope values (G1 arrest time) for the proteins in each abundance quartile. Box plots indicate 5th, 25th, medial, 75th, and 95th percentiles. **(G)** Correlation of protein slopes calculated from the two orthogonal experimental systems. For the left panel, the protein slope calculated from the size mutants used the mean volume measurements (C). For the right panel, the mean volume measurements from (C) were weighted by the relative DNA content shown in (A). The “adjusted” protein slope more accurately reflects changes in the DNA-to-volume ratio and aligns slightly better with protein slopes calculated from G1-arrested cells. The “adjusted” plot is shown in Figure 2. **(H)** Heatmap depicting the relative concentrations of ∼4000 budding yeast proteins in the G1 arrest time experiment shown in (C) and (D). Proteins are ordered by descending protein slope.

**Figure S4 - Supplement for Figure 2.**
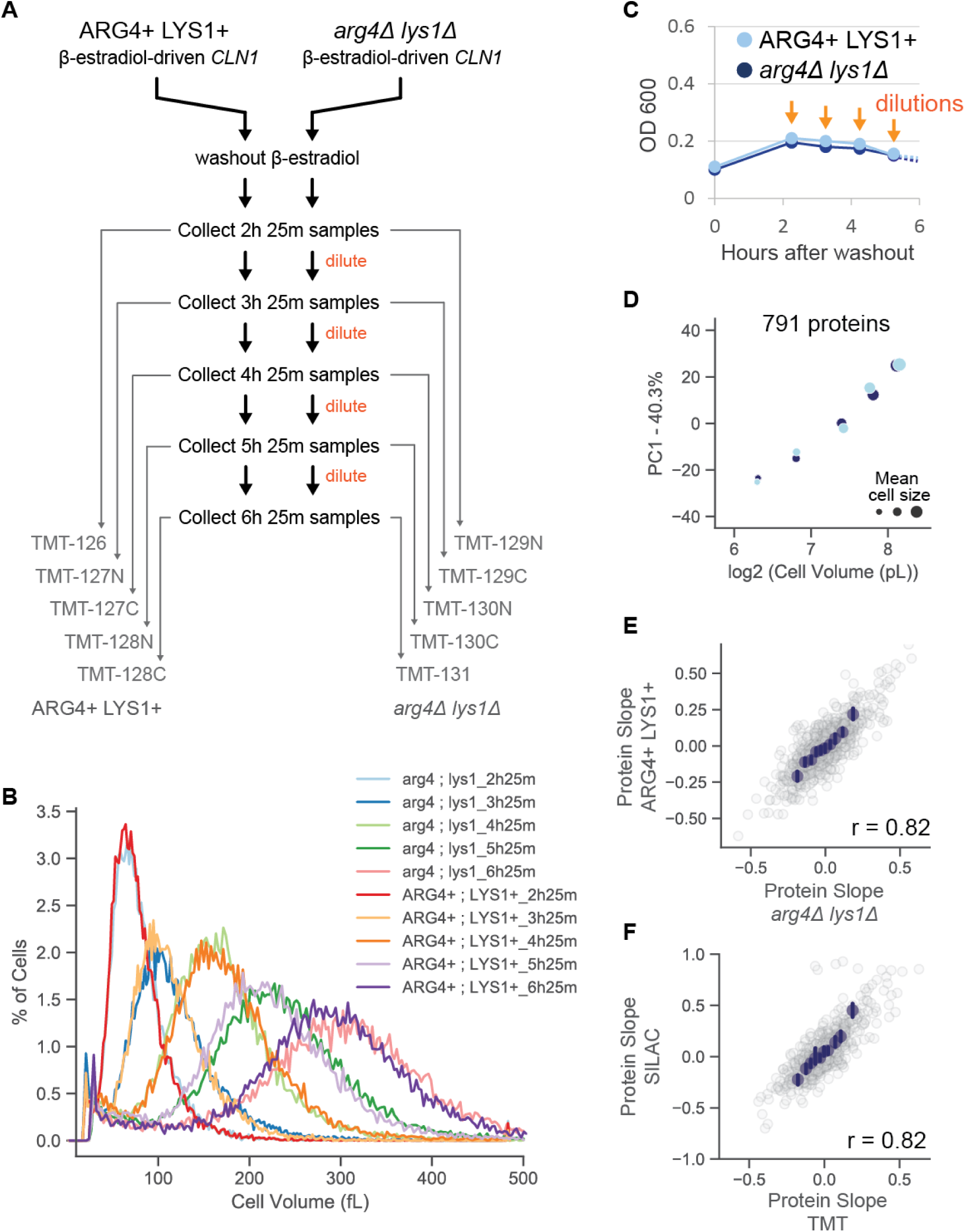
**(A)** Experimental scheme to test whether SILAC-related mutations or culture density affects size-dependent proteome changes. The indicated samples were labeled with 10-plex TMT. A protein slope value was calculated in a manner similar to Figure 2D. Here, TMT MS3 reporter ions were used to quantify relative changes in protein concentration rather than MS1 SILAC ions. **(B)** Cell size distributions determined by a Coulter counter. Time points of the G1 arrest time course are differentially colored. **(C)** During the G1 arrest time course, culture flasks were repeatedly diluted with pre-warmed media to maintain a near constant culture density. **(D)** Principal component analysis of the proteome measurements of the G1 arrested cells. The 1st principal component is plotted against mean cell volume. Dot size represents mean cell volume. Colors represent the replicate experiments in the different strain backgrounds depicted in (A). **(E)** Correlation of Protein Slope values calculated in the *arg4Δlys1Δ* and *ARG4 LYS1* strain backgrounds from (A). Blue dots are x-binned data and error bars represent the 99% confidence interval. r value denotes the Pearson correlation coefficient. **(F)** Correlation of Protein Slope values calculated using TMT with the protein slope measurements calculated using SILAC (G1 arrest time, Figure 2). Blue dots are x-binned data and error bars represent the 99% confidence interval. r value denotes the Pearson correlation coefficient.

**Figure S5 - Supplement for Figure 3.**
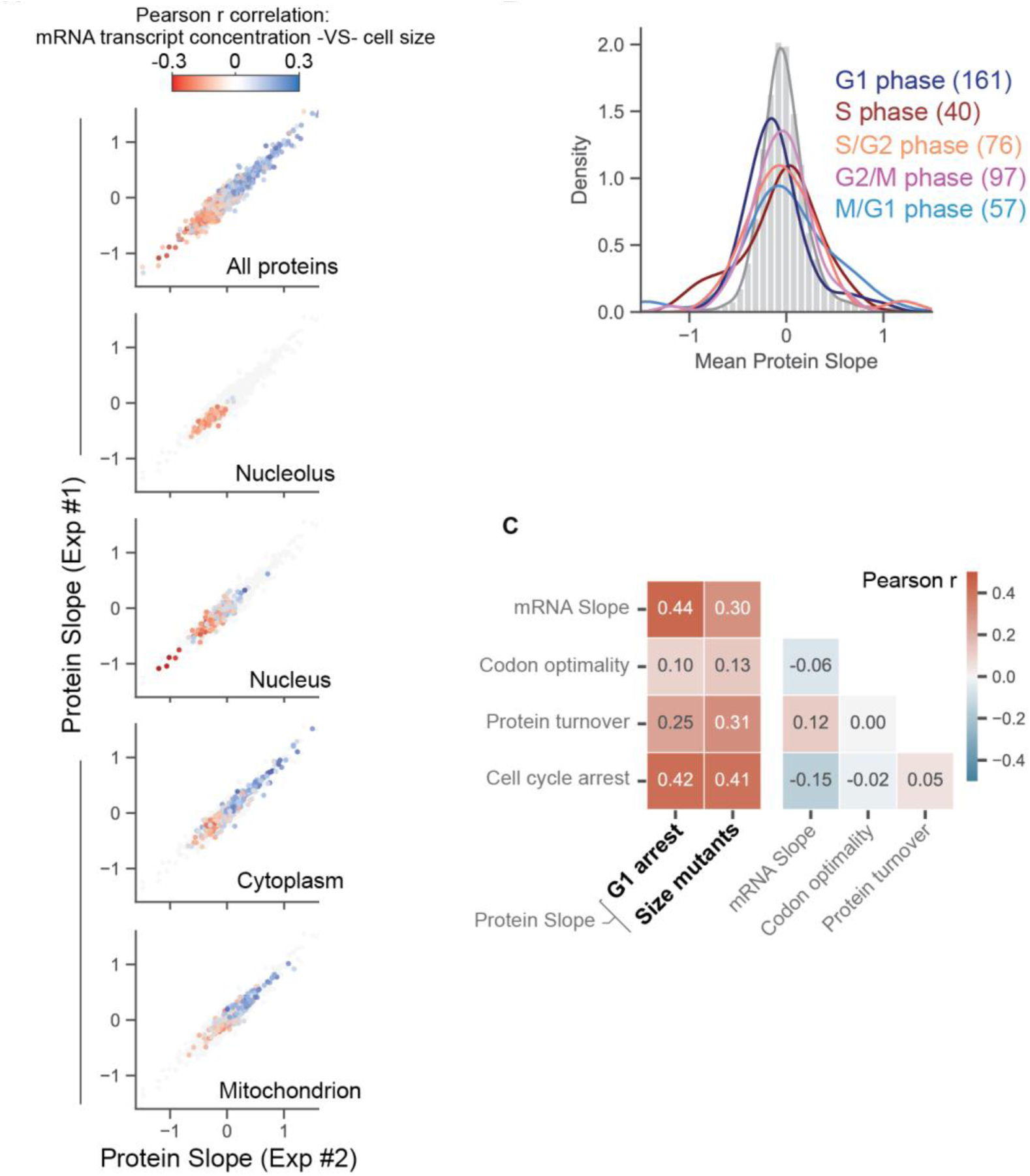
**(A)** Strategy to determine the size scaling behavior of individual transcripts from the mRNA microarray measurement of 1,484 yeast mutant strains. For each measured transcript, its relative concentration in each mutant strain was regressed against the size of that mutant strain. Pearson r values reflect the size-scaling behavior of individual mRNA transcripts. Plots showing the size-scaling behaviors of proteins with subcellular localizations are colored based on the transcript Pearson r value. **(B)** Histograms of protein slope values from the G1 arrest time experiment in Figure 2. Colored lines represent distribution of slope values for proteins whose expression is enriched in the indicated cell cycle phase ^69^. The number of proteins represented in the histogram is in parentheses. Histogram of all protein slope values is shown in gray. **(C)** Pearson correlation matrix for dependent and independent variables in the OLS regression model. Collinearity of independent variables is minimal.

**Figure S6 - Supplement for Figure 4.**
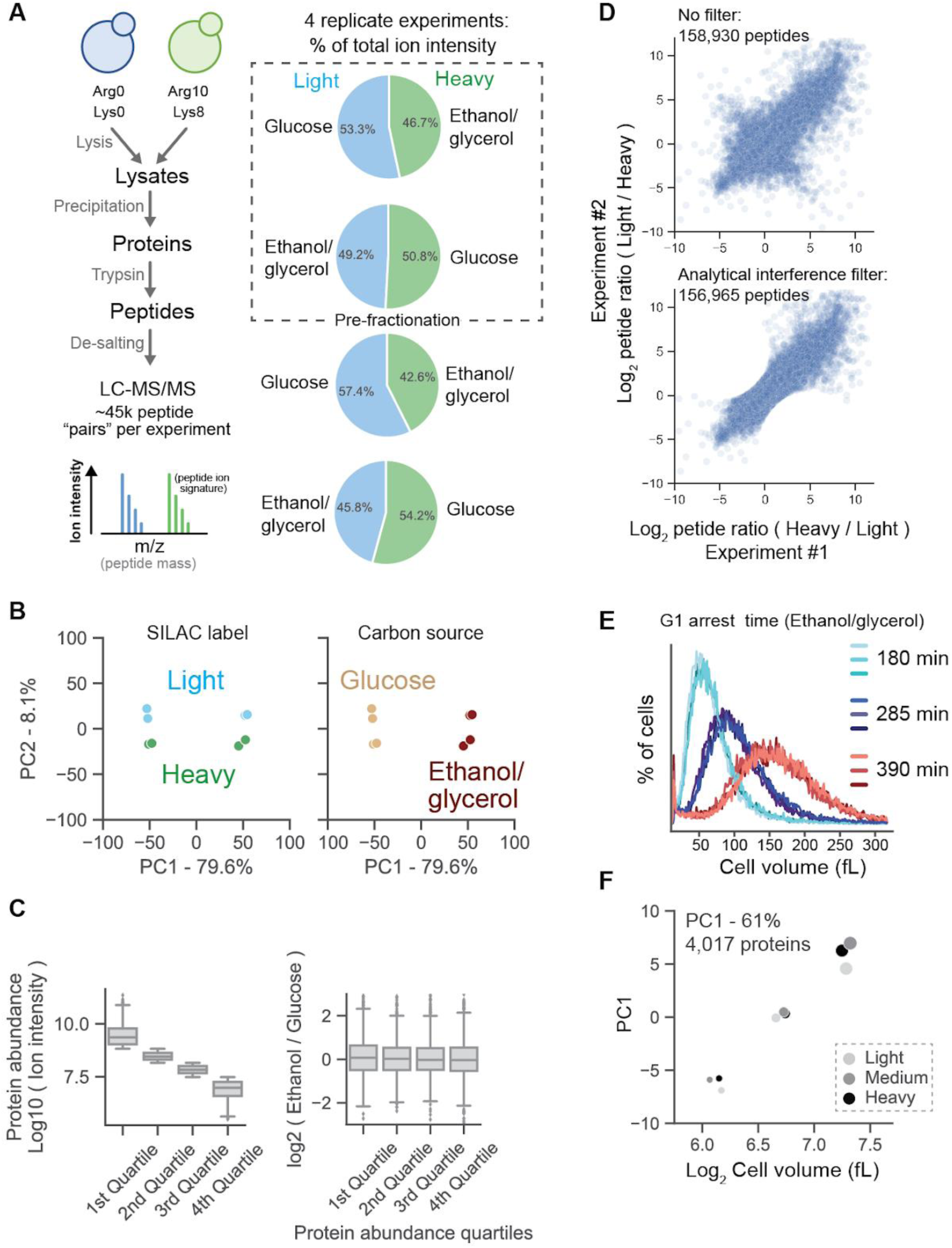
**(A)** Proteome comparison of budding yeast using fermentable and non-fermentable carbon sources. Four SILAC-swapping biological replicate experiments were initially performed. Two of these experiments were then pre-fractionated for deeper proteome analysis. The media mixtures used in the comparison were synthetic complete with 2% dextrose (Glucose) and synthetic complete with 1% ethanol and 2% glycerol (Ethanol/glycerol). **(B)** Principal component analysis of all 4 biological replicate experiments. The same plot is shown twice. Coloring highlights the source of variance for PC1 and PC2. **(C)** For each individual protein, a crude estimation of copy number (summed peptide intensity) was used to bin by abundance quartiles. Protein abundance does not influence the degree to which a protein’s concentration changes between growth conditions. **(D)** Correlation of SILAC ratios for every unique peptide measurement shared between two biological replicate experiments. A unique peptide measurement is defined by the peptide sequence, modification state, charge state, and fraction number. To filter out peptide measurements that were contaminated by analytical interference, peptides that produced a reciprocal measurement between SILAC-swapped replicates were excluded (bottom plot). **(E)** G1 arrest time course (as described in Figure 2) was performed in synthetic complete media supplemented 1% ethanol and 2% glycerol as a carbon source. The length of the G1 arrest time course was extended due to the slower growth rate in ethanol/glycerol media. Size separation was confirmed using a Coulter counter. Light-, medium-, and heavy-labeled yeast are differentially shaded. **(F)** Principal component analysis of the proteome measurements on small-, medium-, and large-sized cells grown in ethanol/glycerol media. The 1st principal component is plotted against mean cell volume. Dot size represents mean cell volume. Colors represent the SILAC label. SILAC labeling orientation for small, medium, and large cells was swapped for replicate experiments. All three replicate experiments are plotted together.

**Figure S7 - Supplement for Figure 5.**
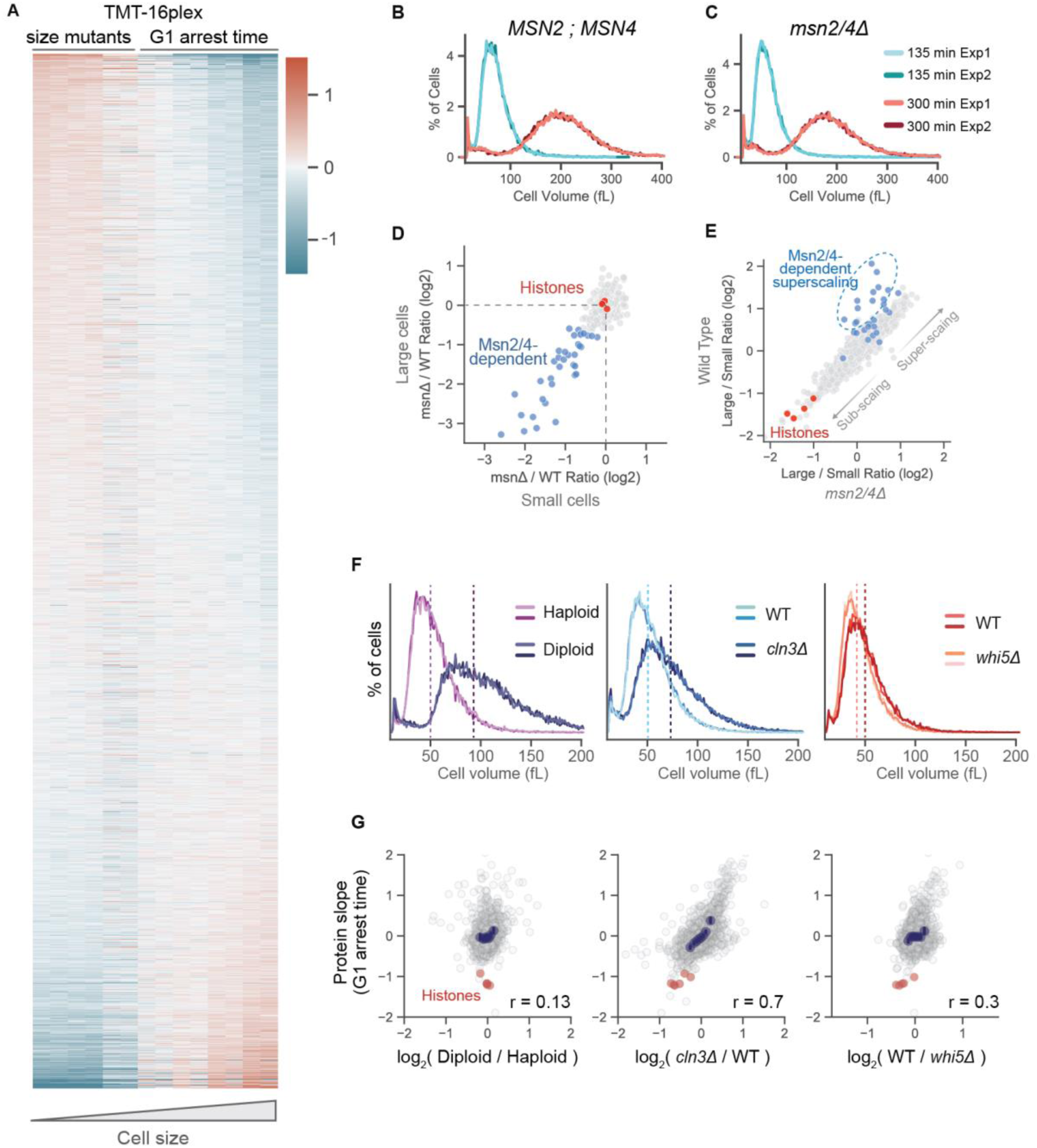
**(A)** Heatmap of relative protein concentration changes across different cell sizes. Each size is represented by two replicate columns. Size mutants correspond to asynchronous *whi5Δ*, WT, and *cln3Δ* cultures. G1 arrest time corresponds to 4 time points after G1 arrest taken at 1-hour intervals (using the genetic systems described in Figure 2). **(B and C)** Coulter counter measurements of (B) *MSN2 MSN4* and (C) *msn2Δmsn4Δ* G1-arrested cells. Biological replicate experiments are differentially shaded. **(D)** Proteins whose expression is dependent on *msn2Δmsn4Δ* are denoted in blue. Proteins highlighted in blue are the same set depicted in Figure 5E. Msn2/4-dependency was defined as decrease in concentration of > 1.6- fold. **(E)** Figure 5E is re-plotted here for reference. The same genes identified in (D) as Msn2/4-dependent are shown in blue. **(F)** Coulter counter measurements of the indicated strains. Biological replicate experiments are differentially shaded. SILAC channels were swapped for replicate experiments to maximize measurement accuracy. **(G)** Correlation of protein slope values (G1 arrest time) with the concentration ratios calculated from the binary comparison of the indicated strains. Each concentration ratio is the average of two SILAC label-swapped replicate experiments. Core histone proteins are shown in red. Blue dots are x-binned data and error bars represent the 99% confidence interval. r value denotes Pearson correlation coefficient.

**Figure S8 - Supplement for Figure 6.**
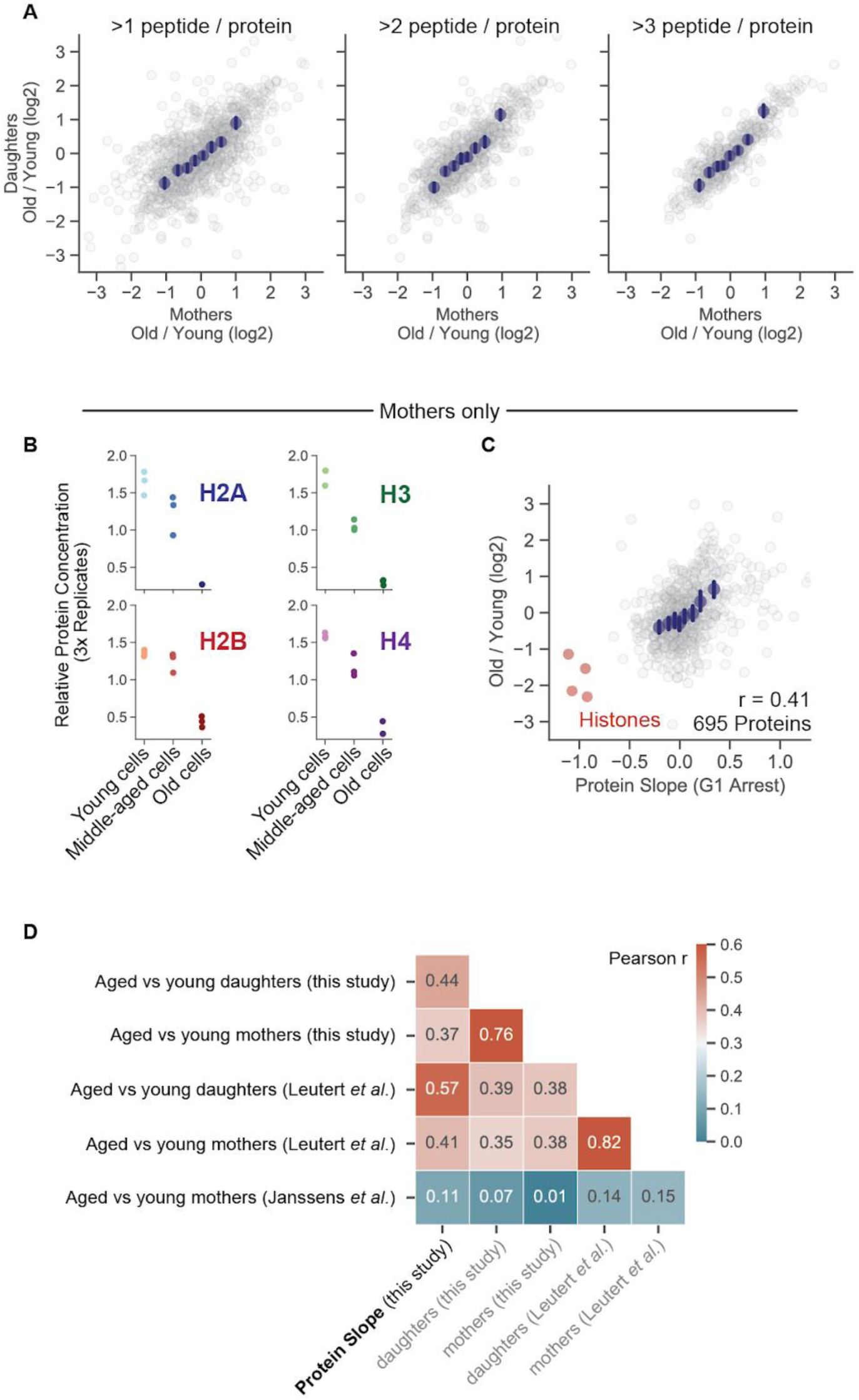
**(A)** Correlation of log_2_ (old / young) concentration ratios derived from young and old budding yeast mothers and their daughter cells. **(B)** Relative concentrations of core histone proteins in young, middle-aged, and old yeast mothers. Each dot corresponds to 1 of 3 biological replicate experiments. **(C)** Correlation of age-associated and size-dependent proteome changes. Log_2_ (old / young) concentration ratio is calculated from mother cells. Protein slope value is from the G1 arrest time experiment in Figure 2. Blue dots are x-binned data and error bars represent the 99% confidence interval. r value denotes Pearson correlation coefficient. **(D)** Correlation grid containing size scaling and aging proteomics data. Protein slope values (Figure 2), log_2_ (old / young) concentration ratio for daughter (Figure 6) and mother (B), and similar measurements from the literature ^43,70^ were cross correlated together. Numbers and colors denote Pearson correlation value for interesting datasets.

## Methods

### Primary hepatocyte isolation, culture, and FACS sorting

Primary hepatocytes from Rosa26-CreER2; FUCCI2 mice were isolated by two-step collagenase perfusion ^71^. Liver perfusion medium (Thermo Fisher Scientific, 17701038), liver digest medium (Thermo Fisher Scientific, 17703034) and hepatocyte wash medium (Thermo Fisher Scientific, 17704024) were used. After isolation, cells were subjected to 2D culture. The protocol for 2D culture of primary hepatocytes was kindly shared by Yinhua Jin from Dr. Roeland Nusse’s lab at Stanford University. More specifically, after isolation, cells were washed 3 times with hepatocyte wash medium (Thermo Fisher Scientific, 17704024). Cells were then plated in a 6-well plate precoated with collagen I (50µg/mL) at a density of 200,000 per well. The culture medium contained 3µM CHIR99021 (Peprotech), 25ng/mL EGF (Peprotech), 50ng/mL HGF (Peprotech), and 100ng/mL TNFa (Peprotech) in Basal medium. The Basal medium contained William’s E medium (GIBCO), 1% Glutamax (GIBCO), 1% Non-Essential Amino Acids (GIBCO), 1% Penicillin/streptomycin (GIBCO), 0.2% normocin (Invitrogen), 2% B27 (GIBCO), 1% N2 supplement (GIBCO), 2% FBS (Corning), 10mM nicotinamide (Sigma), 1.25mM N-acetylcysteine (Sigma), 10µM Y27632 (Peprotech), and 1µM A83-01 (Tocris). The culture medium was refreshed every other day. Cells were passaged via trypsinization using TrypLE (Thermo-Fisher).

For FACS sorting, primary hepatocytes were trypsinized and incubated with Hoechst (Thermo Fisher Scientific, 62249) at 10µg/mL at 37°C for 30 minutes. Then, cells were washed with hepatocyte wash medium and sorted using a Flow Cytometer (BD FACSAria II) based on their FUCCI cell cycle marker and DNA staining (see **Figure S1**). Only G1 (with high mCherry-Cdt FUCCI signal and low Venus-Geminin FUCCI signal) hepatocytes with the same ploidy were sorted. Side-scatter (SSC-A) was used to indicate size ^16^, and 3 size bins were gated to sort small, medium, large cells. 10000-20000 cells were sorted for each population. After sorting, 10% of the sorted cells were used to measure cell size using a Z2 Coulter counter (Beckman), and the rest of the cells were lysed in RIPA lysis buffer supplemented with protease inhibitors and phosphatase inhibitors. Proteins were then denatured, reduced, alkylated, and precipitated as described in the LC-MS/MS sample preparation section.

### Budding yeast growth

All yeast were grown in synthetic complete minimal media (US Biological, Y2030) with standard nitrogen, carbon (2% dextrose), and amino acid supplements unless otherwise indicated. Yeast grown for SILAC analysis were cultured in synthetic minimal media lacking lysine and arginine. These cultures were supplemented with “light”, “intermediate”, or “heavy” versions of Lysine (115µM) and Arginine (200µM) (Cambridge Isotope Laboratories). The “light” (Agr0 ; Lys0) version of the media contained L-Arginine and L-Lysine built with normal 12C and 14N isotopes; the “intermediate” (Arg6 ; Lys4) version had L-Arginine containing six 13C atoms and L-Lysine containing four deuterium atoms; the “heavy” (Arg10 ; Lys8) version had L-Arginine containing six 13C and four 15N atoms and L-Lysine containing six 13C and two 15N atoms. To ensure complete labeling, the cells were grown in SILAC media for approximately ∼10 doublings prior to starting an experiment.

Cell cycle arrest experiments were performed using the strain yML127 (prototrophic) or the strain yML129 (*arg1Δlys4Δ*). These strains require Ꞵ-estradiol (final concentration of 20nM, Sigma E2758) to proliferate. To initiate the G1 arrest time course, Ꞵ-estradiol was removed from the culture by centrifugation followed by a single pre-warmed media wash (no Ꞵ-estradiol in wash media). Alternatively, cells were collected on a 0.2µm disposable filter, then washed with pre-warmed media. Washed cells were then resuspended in media without Ꞵ-estradiol (t = 0 minutes of the arrest time course). After 2 hours and 15 minutes in (-)Ꞵ-estradiol media roughly 95% of the yeast are unbudded (**Figure 2B**). Mean cell size was determined using a Z2 Coulter counter (Beckman).

### Large scale isolation of mothers

The Mother Enrichment Program ^42^ was used to isolate old budding yeast mothers. Strain YDJ 1334 was grown to exponential phase and 6×10^8^ cells were distributed between two sterile 15mL conical tubes. The cells in each tube were then spun down, washed with PBS, resuspended in 500µl PBS, and labeled for 30 min with 3 mg of EZ-Link Sulfo-NHS-LC-Biotin (in dark, gentile agitation, and at room temperature). Biotin labeled cells were washed 3x with 1mL of PBS and then split into 3 aliquots. One aliquot was stored immediately at -80°C (young yeast) and the other two were each used to inoculate 1L of YPD culture, one that grew for 24h (middle-aged mothers) and the other that grew for 48h (old mothers). Ꞵ-estradiol (Sigma E2758) was then added to 1µM in each liquid culture for the duration of the incubation period. Strain YDJ 1334 contains genetic modifications that allow for the selective killing of newly born daughter cells in the presence of Ꞵ-estradiol.

To isolate mothers after the 24- or 48-hour incubation period, each 1L culture was spun down, washed 2x with PBS and resuspended in a 50mL conical tube with 10mL of PBS. Resuspended cells were then mixed with 500 µMACS streptavidin magnetic beads (Miltenyi Biotec, Cat #: 130-074-101) and nutated for 30 minutes (in dark at room temperature). Streptavidin-labeled cells were washed two more times with PBS and then resuspended in 80mLs of PBS. Streptavidin-labeled mothers were then captured on LS+ Positive Selection Columns (Miltenyi Biotec, Cat #: 130-042-401). In brief, 4 columns were used for each timepoint (24 and 48 hours). Each column was placed in a magnet quadro MACS (Miltenyi Biotec) and rinsed with 5mL of PBS. Cells were loaded into each column 8 mLs at a time (20mLs in total for each column) and then washed 2x with 8mL of PBS. Column-bound cells were then eluted by gravity flow (14mL of PBS) after removal from the magnet.

### LC-MS/MS sample preparation - SILAC

See Table S10 for a detailed accounting of all proteomic experiments. Yeast cultures were pelleted by centrifugation at 10,000xg for 5 minutes or collected on a 0.2µm disposable filter. Pellets were resuspended in 300µL yeast lysis buffer (50mM Tris, 150mM NaCl, 5mM EDTA, 0.2% Tergitol, pH 7.5) with 700µl of glass beads. Lysis was performed at 4°C in a Millipore Fastprep24 (settings: 6.0m/s, 4×40s). Cell lysates were cleared by centrifugation at 12,000xg for 5 minutes at 4°C. The mixed lysates were then denatured in 1% SDS, reduced with 10mM DTT, alkylated with 5mM iodoacetamide, and then precipitated with three volumes of a solution containing 50% acetone and 50% ethanol. Proteins were re-solubilized in 2 M urea, 50 mM Tris-HCl, pH 8.0, and 150 mM NaCl, and then digested with TPCK-treated trypsin (50:1) overnight at 37°C. Trifluoroacetic acid and formic acid were added to the digested peptides for a final concentration of 0.2%. Peptides were desalted with a Sep-Pak 50mg C18 column (Waters). The C18 column was conditioned with 500µl of 80% acetonitrile and 0.1% acetic acid and then washed with 1000µl of 0.1% trifluoroacetic acid. After samples were loaded, the column was washed with 2000µl of 0.1% acetic acid followed by elution with 400µl of 80% acetonitrile and 0.1% acetic acid. The elution was dried in a Concentrator at 45°C.

### LC-MS/MS sample preparation - TMT

Lysis, denaturation, reduction, and precipitation for SILAC analysis was the same for TMT analysis (Lysis buffer and working solution of Iodoacetamide used HEPES instead of Tris). Our method for TMT labeling was adapted from Zecha et al. ^72^ and the Thermo TMT10plex™ Isobaric Label Reagent Set Protocol. In brief, acetone precipitated proteins were resuspended in 100µm TEAB and digested overnight with TPCK trypsin (50:1) in the absence of Tris or Urea. After digestion, the peptide concentration was ∼1µg/μL in 100µM TEAB for all samples. 20µg of peptide was labeled using 100µg of Thermo TMT10plex™ or TMTpro™ 16plex in a reaction volume of 25µl for 1 hour. The labeling reaction was quenched with a final concentration of 0.5% hydroxylamine for 15 minutes. Labeled peptides were pooled, acidified to a pH of ∼2 using drops of 10% trifluoroacetic acid, and desalted with a Sep-Pak 50mg C18 column as described above.

### High-pH reverse phase fractionation

SILAC- or TMT-labeled peptides were fractionated using a Pierce™ High pH Reversed-Phase Peptide Fractionation kit. The eight default fractions were either injected separately or pooled back into 4 fractions (1-5, 2-6, 3-7, 4-8). Dried peptides were reconstituted in 0.1% trifluoroacetic acid (TFA). Peptide concentrations were determined using a nanodrop prior to injection.

### LC-MS/MS data acquisition - SILAC

Desalted SILAC-labeled peptides were analyzed on a Fusion Lumos mass spectrometer (Thermo Fisher Scientific, San Jose, CA) equipped with a Thermo EASY-nLC 1200 LC system (Thermo Fisher Scientific, San Jose, CA). Peptides were separated by capillary reverse phase chromatography on a 25 cm column (75 µm inner diameter, packed with 1.6 µm C18 resin, AUR2-25075C18A, Ionopticks, Victoria Australia). Peptides were introduced into the Fusion Lumos mass spectrometer using a 125 min stepped linear gradient at a flow rate of 300 nL/min. The steps of the gradient are as follows: 3–27% buffer B (0.1% (v/v) formic acid in 80% acetonitrile) for 105 min, 27-40% buffer B for 15 min, 40-95% buffer B for 5min, maintain at 90% buffer B for 5 min. Column temperature was maintained at 50°C throughout the procedure. Xcalibur software (Thermo Fisher Scientific) was used for the data acquisition and the instrument was operated in data-dependent mode. Advanced peak detection was enabled. Survey scans were acquired in the Orbitrap mass analyzer (Profile mode) over the range of 375 to 1500 m/z with a mass resolution of 240,000 (at m/z 200). For MS1, the Normalized AGC Target (%) was set at 250 and max injection time was set to “Auto”. Selected ions were fragmented by Higher-energy Collisional Dissociation (HCD) with normalized collision energies set to 31 and the tandem mass spectra was acquired in the Ion trap mass analyzer with the scan rate set to “Turbo”. The isolation window was set to 0.7 m/z window. For MS2, the Normalized AGC Target (%) was set to “Standard” and max injection time was set to “Auto”. Repeated sequencing of peptides was kept to a minimum by dynamic exclusion of the sequenced peptides for 30 seconds. Maximum duty cycle length was set to 1 second.

### LC-MS/MS data acquisition - TMT

Instrumentation and ionization for TMT analysis was the same as the SILAC experiments described above. Peptides were introduced into the Fusion Lumos mass spectrometer using a 180 min stepped linear gradient at a flow rate of 300 nL/min. The steps of the gradient are as follows: 6–33% buffer B (0.1% (v/v) formic acid in 80% acetonitrile) for 145 min, 33-45% buffer B for 15 min, 40-95% buffer B for 5 min, maintain at 90% buffer B for 5 min. Column temperature was maintained at 50°C throughout the procedure. Xcalibur software (Thermo Fisher Scientific) was used for the data acquisition and the instrument was operated in data-dependent mode. Advanced peak detection was disabled. Survey scans were acquired in the Orbitrap mass analyzer (centroid mode) over the range of 380 to 1400 m/z with a mass resolution of 120,000 (at m/z 200). For MS1, the Normalized AGC Target (%) was set at 250 and max injection time was set to 100ms. Selected ions were fragmented by Collision Induced Dissociation (CID) with normalized collision energies of 34 and the tandem mass spectra was acquired in the Ion trap mass analyzer with the scan rate set to “Rapid”. The isolation window was set to 0.7 m/z window. For MS2, the Normalized AGC Target (%) was set to “Standard” and max injection time was set to 35ms. Repeated sequencing of peptides was kept to a minimum by dynamic exclusion of the sequenced peptides for 30 seconds. The maximum duty cycle length was set to 3 seconds. Relative changes in peptide concentration were determined at the MS3-level by isolating and fragmenting the 5 most dominant MS2 ion peaks.

### Spectral searches - TMT and SILAC

All raw files were searched using the Andromeda engine ^73^ embedded in MaxQuant (v2) ^74^. In brief, 3 label SILAC search was conducted using Maxquant’s default Arg6/10 and Lys4/8 labels. For TMT searches, a Reporter ion MS3 search was conducted using 10plex or TMTpro isobaric labels. For both TMT and SILAC searches, variable modifications included oxidation (M) and protein N-terminal acetylation. Carbamidomthyl (C) was a fixed modification. The number of modifications per peptide was capped at five. Digestion was set to tryptic (proline-blocked). For SILAC experiments, peptides were not “Re-quantified”, and maxquant’s match-between-runs feature was not enabled. Database search was conducted using the UniProt proteome - Yeast_UP000002311_559292. The minimum peptide length was 7 amino acids. 1% FDR was determined using a reverse decoy proteome.

### RNA extraction and sequencing

RNA was extracted using from yeast pellets using Direct-zol™ RNA Microprep Kit (Zymo Research). mRNA was enriched using the NEBNext Poly(A) mRNA Magnetic Isolation Module (NEB, #E7490). The NEBNext Ultra II RNA Library Prep Kit for Illumina® (NEB, #E7775) was then used to prepare libraries for paired-end (2×150 bp) Illumina sequencing (Novogene). RNAseq data processing was performed as described previously ^19^.

### Peptide quantitation - SILAC

Our SILAC analysis pipeline was employed previously ^19^. We used the peptide feature information in MaxQuant’s “evidence.txt” output file. Peptide triplets (each row in the “evidence.txt” table) are assigned to a protein based on MaxQuant’s “Leading razor protein” designation. For each peptide triplet, the fraction of ion intensity in each SILAC channel was calculated by dividing the “Intensity L/M/H” column by the “Intensity” column. SILAC channels were normalized by adjusting the fraction of ion intensity in each channel by the median for all measured peptides (see **Figure S2B and S2C**). After normalization, the relative signal difference between the SILAC channels for each peptide triplet was plotted against the normalized cell size (**Figure 1B**). A unique peptide measurement is defined by the peptide sequence, modification state, charge state, and fraction number.

For each unique peptide measurement, we calculated its slope as follows:

Y_1,2,3_ = Relative signal in each SILAC channel (order based on labeling orientation)

Avg. size = (mean volume of small bin + mean volume of medium bin + mean volume of large bin) / 3

x_1_ = (Mean volume of small size bin) / Avg. size

x_2_ = (Mean volume of medium size bin) / Avg. size

x_3_ = (Mean volume of large size bin) / Avg. size

Based on the expectation that our experimental conditions would not result in large, non-linear changes in protein expression, we excluded peptide triplets whose three data points did not loosely fit a linear regression line. Linear regressions on the ∼50,000 triplets/experiment were performed using np.polyfit in Python. Regressions with a mean squared error > 0.075 were excluded. Because this filtering step significantly improved the overall data quality (**Figure S2E**), we concluded that our filtering method mostly excludes peptide triplets contaminated by analytical interference or near the noise floor.

Individual peptide measurements were consolidated into a protein level measurement using Python’s groupby.median. Peptides with the same amino acid sequence that were identified as different charge states or in different fractions were considered independent measurements. We summarize the size scaling behavior of individual proteins as a slope value derived from a regression (similar to what is described above for individual peptides), and each protein slope value is based on the behavior of all detected peptides.

For a given protein, we calculate its slope as follows:

y_i_ = Relative signal in the i^th^ SILAC channel (median of all peptides measured for this protein)

x_i_ = same normalized cell size x_i_ as for the peptide slope calculations above

The protein slope value was determined from a linear fit to the log-transformed data using the equation:

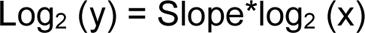

Variables were log-transformed so that a slope of 1 corresponds to an increase in protein concentration that is proportional to the increase in volume and a slope of -1 corresponds to 1/volume dilution. Pearson r and p values for correlation analyses were calculated using scipy’s pearsonr module in Python.

### Peptide quantitation - TMT

Our TMT analysis pipeline uses the peptide feature information in MaxQuant’s “evidence.txt” output file. Each row of the “evidence.txt” file represents an independent peptide and its corresponding MS3 TMT measurement. Peptides without signal in any of the TMT channels were excluded. TMT peptide measurements were assigned to protein based on MaxQuant’s “Leading razor protein” designation. For each peptide measurement, the fraction of ion intensity in each TMT channel was calculated by dividing the “Reporter ion intensity” column by the sum of all reporter ion intensities.

TMT channels were normalized by adjusting the fraction of ion intensity in each channel by the median for all measured peptides (similar to the SILAC normalization in **Figure S2B,C**). After normalization, the relative signal difference between the TMT channels for each peptide triplet was plotted against the normalized cell size for each of the bins of isolated G1 cells. Slope values in Figure 2F and 2G were derived in a manner analogous to the Slope values calculated in the SILAC experiments. Pearson r and p values for correlation analyses were calculated using SciPy’s pearsonr module in Python.

### Orthology analysis

Human ortholog pairs were retrieved using the DRSC Integrative Ortholog Prediction Tool (DIOPT) found at https://www.flyrnai.org/cgi-bin/DRSC_orthologs.pl ^27^. Yeast or mouse proteins were used as the “input” species and humans were set as the “output”. For budding yeast, the systematic gene name was used for the “Search term” and, for mice, the uniprot ID was used. We only considered ortholog pairs with a DIOPT “weighted Score” of greater than 10 for our analyses. Once mouse or yeast proteins were matched with a human ortholog protein, we imported protein slope values derived from a human RPE-1 cell line ^19^.

### 2D Annotation Enrichment analysis

Annotation enrichment analysis was performed as described previously ^31^. The protein annotation groups were deemed significantly enriched and plotted if the Benjamini-Hochberg FDR was smaller than 0.02. The position of each annotation group on the plot is determined by the enrichment score (S). The enrichment score is calculated from the rank ordered distribution of Protein Slope values:

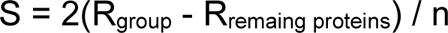

Where R_group_ and R_remaing proteins_ are the average ranks for the proteins within an annotation group and all remaining proteins in the experiment, respectively, and n is the total number of proteins. Highlighted annotation groups were manually curated. All significantly super- and sub-scaling annotation groups can be found in Table S4.

### Principal component analysis

PCA analysis was performed in Python using the sklearn package. A data frame was created that contained individual proteins as rows with columns corresponding to the relative protein concentration in each SILAC or TMT channel (obtained from the median of all peptide measurements for a given protein).

### Protein annotations

For 2D annotation enrichment plots, the default GOMC, GOMF, and GOCC annotations were used. For all other plots, protein localization annotations were based on UniProt columns named “Subcellular location [CC]” or “Protein names”. Protein localization was strictly parsed so that each annotated protein belongs to only one of the designated groups. Proteins with two or more of the depicted annotations were ignored (except for the “Cytoplasm / Nucleus” category, which required both a nuclear and cytoplasmic annotation). ESR- and growth rate-associated proteins were annotated based on previous work ^38^.

### OLS linear regression model

Multiple linear regression analyses were performed using the statsmodels module in Python. The prediction of size scaling behavior was based on the proteins shared between the protein turnover (villen), mRNA Slope, and Protein Slope datasets (at least 2 peptides / protein). Independent variables for codon score (YeastMine), mRNA Slope, and protein turnover (T50%) were each independently standardized by subtracting all values by the dataset’s mean and then dividing by the dataset’s standard deviation. The subcellular localization variable was based on UniProt’s “Subcellular location [CC]” annotations and entered as a binary value for each compartment (1 if a protein possessed an annotation and 0 if it did not). Only subcellular compartments that provided nonredundant predictive power were ultimately included in the model. A constant value was added to the regression equation using the add_constant function in statsmodels. We set the benchmark for predictive accuracy (Prediction %) as the correlation between biological replicate experiments (Protein Slope from Exp #1 vs the average slope from Exp #2 and Exp #3).

